# Ice drives the emergence of functional protocells

**DOI:** 10.64898/2026.10.13.758973

**Authors:** Tatsuya Shinoda, Kimberly Poppy Sinclair, Natsumi Noda, Shunto Harada, Hye-Eun Lee, Ryuhei Nakamura, Tomoaki Matsuura

## Abstract

A defining feature of living systems is the high local concentration of diverse molecular components within membrane-bounded compartments. How did such an organization arise *de novo*, without complex molecular machinery or pre-existing cellular structures that could template their own reproduction? Prebiotic environments may have provided the physical driving forces needed to bring dispersed molecular components together and organize them into functional compartments. Here, we show that a freeze–thaw (FT) process can couple molecular concentration, the formation of cell-sized membrane compartments, and the activation of a biochemical system. *In situ* imaging showed that chemically diverse solutes and membrane material were co-concentrated within ice-confined liquid regions during slow freezing. Subsequent slow thawing gave rise to cell-sized giant vesicles that capture the concentrated molecules. Crucially, we show that these sequential processes can assemble a multicomponent transcription–translation system (PURE system) within vesicle compartments, enabling protein synthesis from diluted mixtures with little or no detectable activity in bulk solution. By demonstrating that functional cellular organization can emerge from otherwise inactive molecular components, our study suggests that environmental FT processes provide a physical route for the transition from chemical to biological organization, with implications for the emergence of life on Earth and beyond.

## Introduction

Living cells are crowded, membrane-bounded systems in which diverse molecular components interact in a coordinated manner^1,2^. How such cellular organization first emerged from prebiotic chemistry remains an enduring mystery. Prebiotic chemistry has established plausible routes to many of life’s molecular building blocks^3–5^, yet in dilute environments, diffusion tends to disperse these molecules. Such dilution reduces molecular encounter rates and slows reaction kinetics, potentially rendering possible chemistry inaccessible on relevant timescales^6,7^. In addition, dilution limits the local availability of amphiphiles required to form membrane compartments, which protects reaction networks from dilution and helps maintain an internal composition distinct from the surroundings^1,8,9^. The emergence of cellular organization therefore requires mechanisms capable of concentrating diverse biological molecules against dilution under prebiotically plausible conditions.

Geophysical processes are known to drive entropically disfavored molecular concentration by coupling it to thermodynamically downhill processes such as heat flow, evaporation and phase transitions. Diverse prebiotic settings have been proposed to exploit such physical driving forces, including intermittently drying hydrothermal pools^10–14^, geothermally heated rock pores^15–21^, freezing–thawing ice environments^22–27^ and mineral surfaces^28,29^, where molecular building blocks can accumulate and simple chemical reactions can be promoted. Yet, these studies have largely focused on the concentration of one or a few molecular species, with limited attention to their encapsulation within membrane-bounded microcompartments. How environmental processes could simultaneously concentrate many distinct molecular components and drive their *de novo* organization into a microcompartment, one of the universal features of all cellular systems, has yet to be demonstrated.

Among the geophysical settings described above, freeze–thaw (FT) processes, which could have occurred on the early Earth^30,31^ and on icy planetary bodies^32^, are of particular interest for the assembly of multicomponent biochemical systems because they can concentrate molecules without requiring extreme heating or severe dehydration. Moreover, unlike molecularly selective mineral adsorption^33^ or thermophoretic transport^19^, freezing can concentrate a broad range of chemically diverse molecules through their exclusion from growing ice.

In this study, we investigated whether FT could couple molecular concentration to the emergence of functional microcompartments. When biomolecules such as proteins or DNA were mixed with small, *empty* lipid particles (~100 nm diameter) and subjected to FT, we found the emergence of giant vesicles (GVs) with micrometer-wide diameters whose interiors were enriched in proteins and DNA. These FT events were observed *in sit*u using fluorescence microscopy equipped with a cooling stage, which provided direct visual evidence linking freeze-induced molecular concentration to the subsequent formation of GVs. Finally, the same FT was applied to a multicomponent biochemical system, the PURE system^34,35^, which is a reconstituted *in vitro* transcription–translation system. The PURE system, which consists of around 70 different molecules, was diluted to the extent that it can no longer synthesize proteins, was mixed with small lipid particles, and was subjected to FT. We observed the emergence of GVs with active PURE system encapsulated inside, demonstrating the assembly of cell-like structure from a diluted and inactive molecular system. Our study suggests that environmental FT processes may provide a physical route for the transition from dilute prebiotic chemistry to functional protocellular organization, with implications for the emergence of life on Earth, and potentially on extraterrestrial worlds.

## Results and discussion

### Slow FT generates giant vesicles and encapsulates small molecules

We initially investigated how nanoscale phospholipid vesicles respond to FT by monitoring changes in vesicle size and molecular uptake. We prepared empty vesicles with an average diameter of 100 nm from three phospholipids with distinct acyl-chain structures: 1-palmitoyl-2-oleoyl-glycero-3-phosphocholine (POPC, 16:0–18:1 PC), 1-palmitoyl-2-linoleoyl-sn-glycero-3-phosphocholine (PLPC, 16:0–18:2 PC) and 1,2-dioleoyl-sn-glycero-3-phosphocholine (DOPC, 18:1–18:1 PC) (Fig. S1). Vesicles suspended in 50 mM HEPES buffer (pH 7.4) were mixed with calcein, a hydrophilic fluorescent small molecule, and subjected to FT treatment with different freezing and thawing rates (Table 1). After the FT treatment, changes in vesicle size were assessed by dynamic light scattering (DLS), while the fraction of encapsulated calcein was quantified after quenching external fluorescence by cobalt ions (Co^2+^). The FT conditions are denoted as F/F for fast freezing and fast thawing, S/S for slow freezing and slow thawing, and similarly for the remaining seven conditions (Table 1).

**Table 1:**
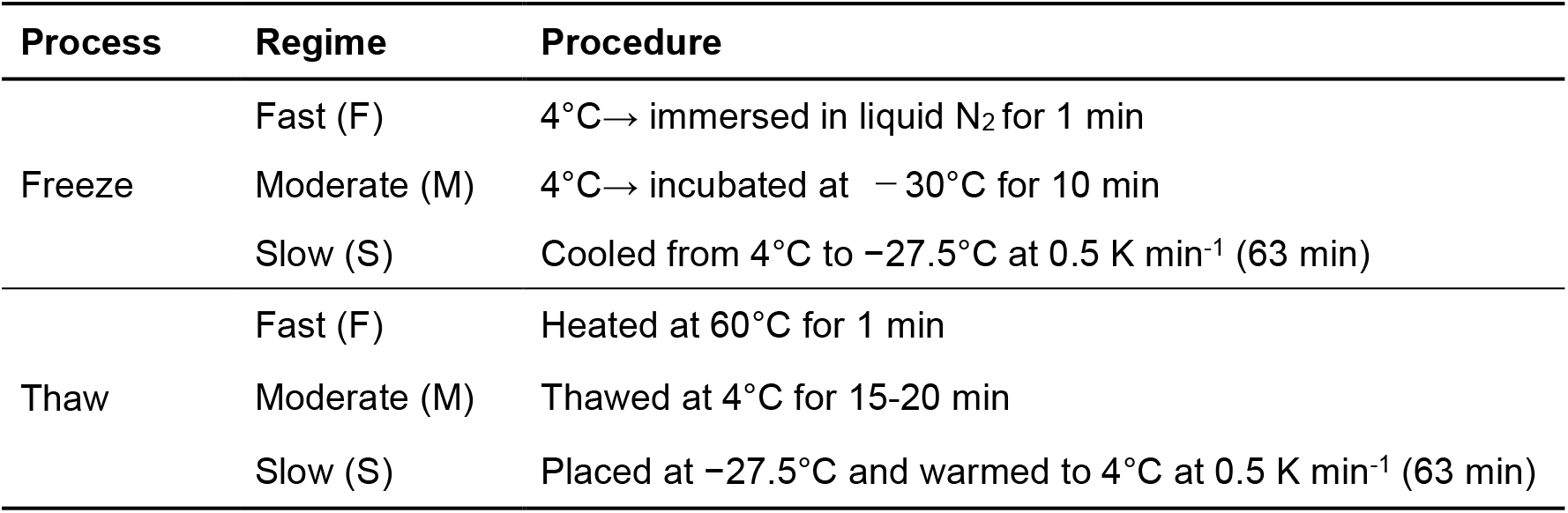
FT conditions used in the current study.

| Process | Regime | Procedure |
| --- | --- | --- |
| Freeze | Fast (F) | 4°C → immersed in liquid N<sub>2</sub> for 1 min |
|  | Moderate (M) | 4°C → incubated at –30°C for 10 min |
|  | Slow (S) | Cooled from 4°C to –27.5°C at 0.5 K min<sup>–1</sup> (63 min) |
| Thaw | Fast (F) | Heated at 60°C for 1 min |
|  | Moderate (M) | Thawed at 4°C for 15–20 min |
|  | Slow (S) | Placed at –27.5°C and warmed to 4°C at 0.5 K min<sup>–1</sup> (63 min) |

DLS revealed the appearance of a larger-vesicle population for PLPC and DOPC under FT conditions that included a slow step (slow freezing or slow thawing), whereas little size increase was observed for POPC (Fig. S2a). Calcein encapsulation showed a similar trend, with higher values under conditions that included a slow step than under those without one; the highest values were observed for DOPC, followed by PLPC and POPC (Fig. 1a).

**Figure 1.**
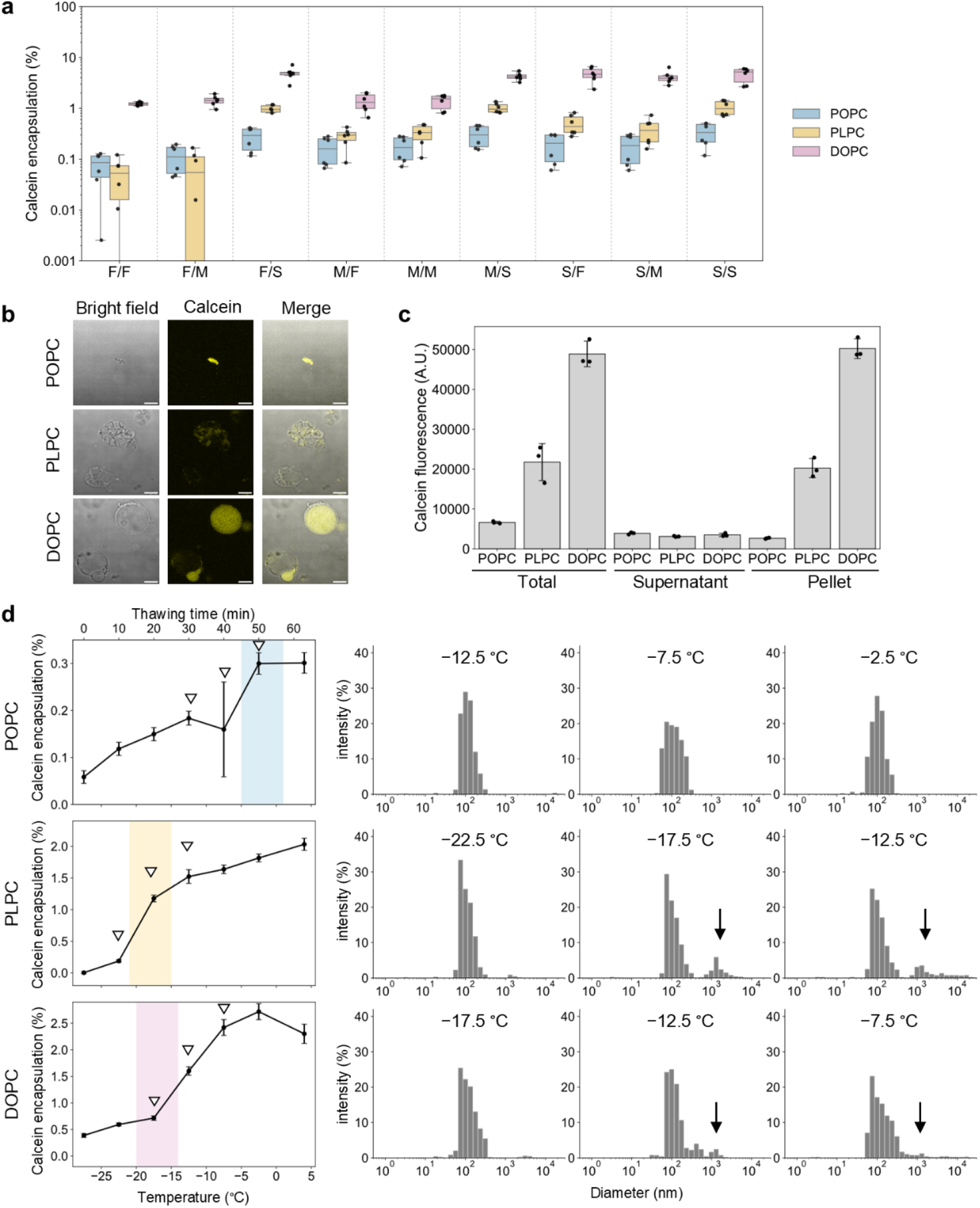
Slow FT enhances vesicle growth and simultaneous small molecule encapsulation. (a) Calcein encapsulation measured across nine FT conditions for vesicles composed of POPC, PLPC, and DOPC. F, M, and S denote fast, moderate and slow, respectively; the first and second letters indicate the freezing and thawing rates, respectively. Calcein encapsulation was calculated as membrane-protected calcein fluorescence after cobalt quenching, normalized to the total fluorescence of calcein initially added to each sample, with both values corrected by subtraction of the corresponding negative-control signals (see Methods). Raw fluorescence data are shown in Supplementary Fig. S6. Box plots show the median and interquartile range, and individual data points represent independent experiments (*n=*6). (b) Representative confocal fluorescence image of giant vesicles (GVs) formed after slow FT (S/S) and cobalt quenching. Scale bar: 10 µm. (c) Calcein fluorescence in the supernatant (non-enlarged vesicle-rich) and pellet (GV-rich) fractions separated by centrifugation after slow FT, and the total fluorescence are shown. Mean ± s.d. are shown (*n=*3). (d) Temperature-resolved analysis of calcein encapsulation and vesicle growth during slow thawing. Left, calcein encapsulation during slow thawing up to the indicated temperature, with the corresponding thawing times shown on the upper axis. Mean ± s.d. are shown (*n=*3). Open triangles indicate the temperatures selected for DLS measurements. The phase transition temperatures (T_m_) of POPC, PLPC and DOPC are −2, −18 and −17 °C, respectively. T_m_ ±3 °C is shaded in color as a visual guide. Right, size distributions measured at the corresponding temperatures; arrows in the DLS plots indicate the fractions of vesicles with increased size.

Under the S/S condition, DOPC vesicles encapsulated 4.60 ± 1.52% of the input calcein, exceeding the theoretical value of 0.36% calculated from the total internal aqueous volume of uniformly dispersed 100 nm vesicles (Methods). This excess encapsulation could arise from molecular enrichment within vesicles (examined further in Fig. 2), the reorganization of many nanoscale vesicles into fewer, larger spherical compartments that would increase the total enclosed aqueous volume available for calcein encapsulation, or both. We first investigated the latter possibility by confocal microscopy and observed the emergence of spherical, micrometer-scale giant vesicles (GVs), either as isolated vesicles or in aggregates, after S/S (Fig. 1b). GVs were scarcely formed by F/F, while S/S gave a significantly larger amount of GVs (Fig. S2b, c). Notably, some of these GVs were unilamellar, as assessed by the α-hemolysin permeability assay^36^ (Fig. S3). To determine whether GV formation accounted for the enhanced calcein encapsulation, we separated GVs from non-enlarged vesicles by standard centrifugation (20,000 × g for 25 min), which predominantly recovered GVs in the pellet fraction while leaving non-enlarged vesicles in the supernatant fraction^37^, and quantified the calcein fluorescence in each fraction. Calcein was highly recovered in the pellet fraction (Fig. 1c), indicating that GVs formed under S/S were the major contributors to calcein encapsulation.

**Figure 2.**
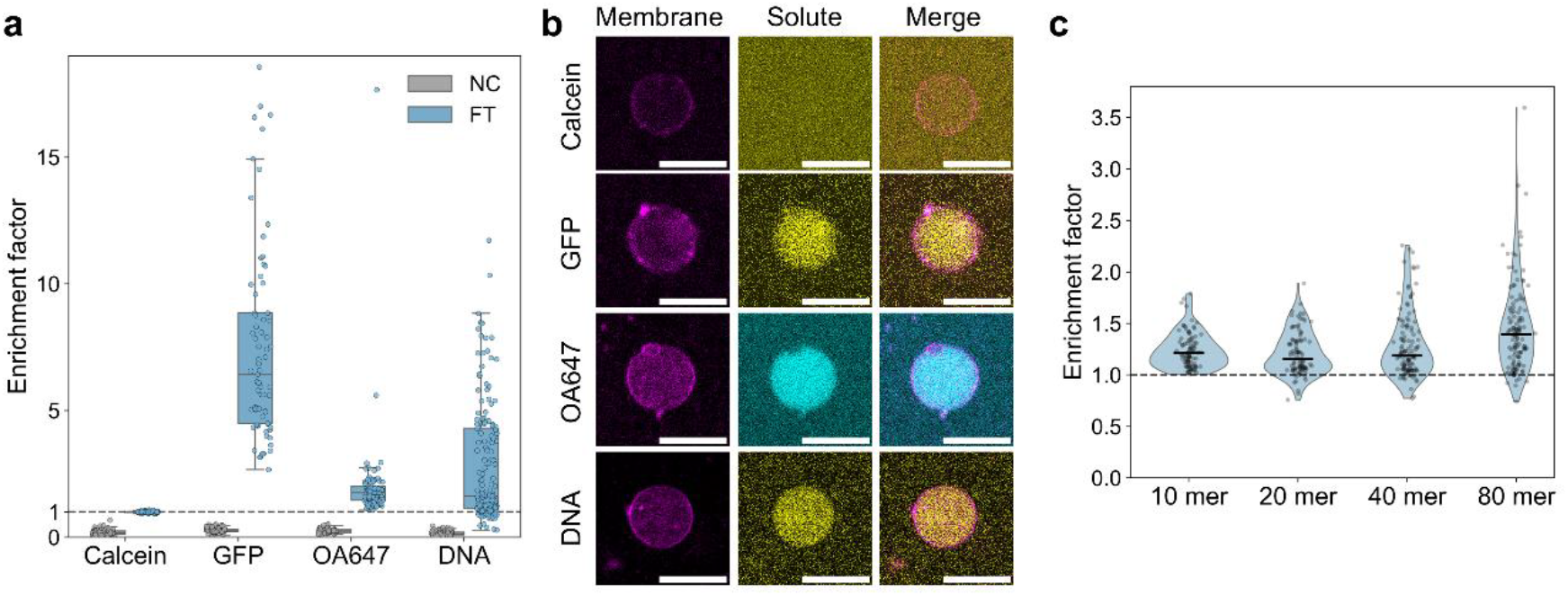
Slow FT enriches diverse biomolecules within newly formed GVs. (a) Enrichment factors of model molecules (calcein, GFP, OA647, and DNA) within DOPC GVs after slow FT. NC denotes the negative control condition, in which molecules were added after slow FT. Box plots show the median and interquartile range, and individual data points represent single vesicles. (b) Representative confocal fluorescence images of DOPC GVs containing model molecules after slow FT. Scale bar: 10 µm. (c) Violin plots of enrichment factors for ssDNA of different lengths within DOPC GVs after slow FT, with horizontal lines indicating the median and individual points representing single vesicles. Summary statistics of enrichment factors are provided in Table S1. See Fig. S8 and Table S2 for further analysis of the enrichment factor distributions.

GV formation depended not only on FT conditions but also on lipid composition. Quantification of the GV fraction, defined as the fraction of total lipid recovered in the GV-enriched pellet after centrifugation, showed that PLPC and DOPC produced substantially higher fractions of GVs than POPC (Fig. S4). Compared with POPC, the presence of more unsaturated bonds in the acyl-chains of PLPC and DOPC is expected to increase membrane disorder and conformational freedom^38,39^, which may facilitate lipid rearrangement at vesicle–vesicle contacts, thereby promoting vesicle fusion and simultaneous solute uptake. Conversely, the lower membrane disorder of POPC may limit such rearrangements, consistent with its limited vesicle growth and calcein encapsulation. Despite comparable GV fractions in PLPC and DOPC, calcein encapsulation was more than two-fold higher in DOPC. To understand the difference between PLPC and DOPC, confocal images during the addition of Co^2+^ were obtained. Upon the addition of Co^2+^, many PLPC GVs lost their encapsulated calcein signal, suggesting a membrane defect. By contrast, many DOPC GVs retained calcein fluorescence, and thus were found to be intact (Fig. S5).

To resolve the dynamics underlying FT-induced vesicle growth and molecular encapsulation, we examined the temperature at which these processes occur during FT. Vesicle suspensions were first fast-frozen and then thawed slowly at 0.5 K min^-1^ in the temperature-controlled bath. At defined temperatures, aliquots were removed from the bath and rapidly thawed. Because fast thawing alone induced little calcein encapsulation (Fig. 1a), the calcein signal measured at each point was attributed primarily to encapsulation that had occurred during slow thawing up to that temperature. Concurrent calcein encapsulation and vesicle growth were observed near the lipid phase transition temperature (Fig. 1d), where lipid membranes exhibit enhanced fluctuations and transient packing defects^40,41^.

Together, these results suggest that S/S promotes the concentration and prolonged contact of vesicles, allowing membrane remodeling to drive vesicle growth and efficient encapsulation of surrounding calcein, predominantly near the lipid phase transition temperature. Among the lipids tested, DOPC most effectively combined vesicle growth with solute retention, yielding spherical micron-scale GVs from nanoscale empty vesicles. We therefore used DOPC vesicles under S/S conditions, hereafter termed slow FT, in subsequent experiments.

### Slow FT enriches biomolecules within the vesicles

During freezing, solutes are expected to concentrate within the interstitial liquid regions between growing ice domains. We therefore asked whether these concentrated solutes could be encapsulated during GV formation. Empty 100 nm DOPC vesicles were mixed with small molecules, proteins, and DNA – namely, calcein (623 Da), SNAP-tagged GFP (48 kDa, hereafter GFP), ovalbumin Alexa Fluor 647 (45 kDa, hereafter OA647), and 956 bp DNA (~591 kDa, SYBR Gold-stained) – and then subjected to slow FT treatment. Again, GV formation was observed, and the molecular enrichment in each individual GV (enrichment factor = *F*_*in*_/ *F*_*out*_) was quantified by comparing the fluorescence inside (*F*_*in*_) versus outside (*F*_*out*_) the vesicles (Fig. 2a, b and Fig. S7). An enrichment factor greater than 1 indicates molecular enrichment within the vesicle, whereas a value of 1 indicates no enrichment. We found that the enrichment factor depended strongly on molecular species. Calcein showed almost no enrichment (median = 0.99-fold), whereas proteins and DNA were enriched inside GVs. The median enrichment factors for GFP, OA647, and DNA were 6.43-, 1.76- and 1.62-fold, respectively (Fig. 2a and Table S1). GFP showed the strongest enrichment on average, with enrichment factor reaching up to 18.6 in a specific GV. When fluorescent molecules were added after GV formation, little uptake was observed, with enrichment factors far below 1, indicating that the resulting GVs were largely impermeable to the tested molecules (Fig. 2a and Table S1).

We next examined the effect of molecular size on enrichment factors. Single-stranded DNA (ssDNA) of different lengths (10, 20, 40, and 80 nt) was subjected to FT. The median enrichment factor was similar for 10-, 20-, and 40-mer ssDNA (1.21-, 1.16-, and 1.19-fold, respectively), but increased to 1.39-fold for the 80-mer, suggesting a moderate size dependence in overall enrichment (Fig. 2c and Table S1). When the distributions of enrichment factors were analyzed for each DNA length, longer DNA showed progressively broader distributions that were increasingly skewed towards high enrichment factors (Fig. 2c, Fig. S8 and Table S2).

The internal molecular composition of FT-generated GVs therefore reflects both molecule-dependent enrichment and substantial vesicle-to-vesicle heterogeneity. The mechanisms giving rise to this heterogeneity, however, remain to be elucidated.

### Slow FT concentrates solutes in ice-confined liquid regions

We next performed *in situ* imaging to directly observe ice formation and the spatial distribution of vesicles and solutes during the FT process. A suspension of rhodamine-labelled DOPC vesicles was placed on a glass slide mounted on a cooling stage, subjected to slow FT, and monitored by time-lapse bright-field and fluorescence imaging (Fig. 3a). The temperature profile used for the *in situ* observation is shown in Fig. 3b. Before ice formation, rhodamine-labelled vesicles were uniformly dispersed. As ice crystals appeared at about −15 °C (38 min), the fluorescence became spatially heterogeneous, forming rhodamine-rich regions. During thawing, ice began to melt at around −2.5 °C (113 min), and the fluorescence turned homogeneous (Fig. 3a). At the same time, rhodamine-rich micron-sized regions appeared (Fig. 3a, inset, arrow), suggesting the formation of GVs, although the imaging resolution did not allow us to unambiguously distinguish spherical vesicles from aggregates.

**Figure 3.**
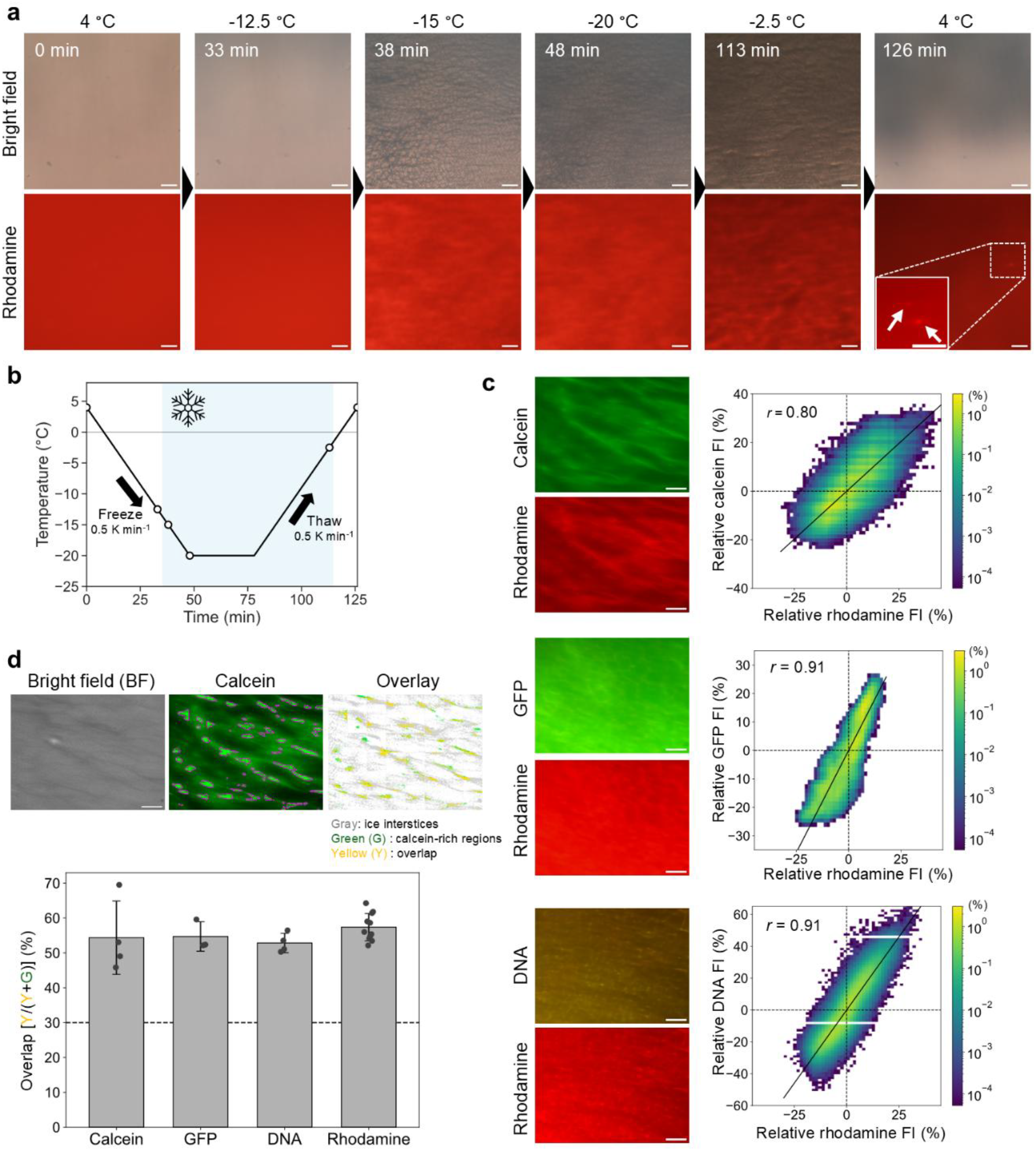
*In situ* observation of ice formation and the spatial distribution of vesicles and solutes during the FT process. (a) Time-lapse bright-field and fluorescence images of rhodamine-labelled DOPC vesicles during the FT process on a cooling stage. Ice crystals formed during cooling and were accompanied by the appearance of rhodamine-rich regions. Upon thawing, fluorescence became more homogeneously distributed across the sample, while rhodamine-rich micron-sized objects remained observable (inset, arrow). Scale bar: 100 µm. (b) Temperature profile used for *in situ* imaging. The sample was cooled from 4 °C to −20 °C at 0.5 K min^-1^, held at −20 °C for 30 min and then warmed at 0.5 K min^-1^. Although the experiments in Figs. 1 and 2 were cooled to −27.5 °C, we limited the *in situ* imaging experiments to −20 °C to operate within the controllable temperature range of the cooling stage. (c) Pixel-wise correlations between vesicle rhodamine fluorescence and solute fluorescence in the frozen state. Relative fluorescence intensity (FI) was calculated as FI = ((*I*− *I*_*mean*_)/*I*_*mean*_) × 100 (%), where *I* is the local pixel fluorescence intensity and *I*_*mean*_ is the mean fluorescence intensity of the entire field. Two-dimensional density plots show relative rhodamine FI versus solute FI, with color indicating pixel frequency. Solid lines indicate linear regression fits, and Pearson’s correlation coefficients are shown in each plot. Scale bar: 100 µm. (d) Spatial overlap of fluorescence-rich regions with dark regions in the corresponding bright-field images. To distinguish local concentration patterns from gradual field-wide intensity variations, each image was divided into 6 × 8 tiles. Within each tile, fluorescence-rich pixels and dark bright-field pixels were defined as the brightest 5% and darkest 30% of pixels, respectively. Fluorescence-rich pixels are highlighted in magenta in the representative calcein fluorescence images. In the overlays, dark bright-field regions are shown in gray, non-overlapping fluorescence-rich pixels in green, and overlapping pixels in yellow. Overlap was calculated as the percentage of fluorescence-rich pixels located within dark bright-field regions (yellow/(green + yellow) × 100 (%)). Bar plots show the mean overlap for calcein, GFP, DNA, and rhodamine, with individual measurements shown as points. The dashed line indicates the overlap expected for a spatially random distribution (30%). Preferential localization of fluorescence-rich regions within dark bright-field regions was consistently observed across different percentile thresholds used to define fluorescence-rich and dark regions. See Fig. S10 for details.

We then examined how model solutes, including calcein, GFP, and DNA, behave when subjected to slow FT together with vesicles. Rhodamine-labelled vesicles were mixed with calcein, GFP, or DNA and imaged *in situ* after slow freezing to −20 °C. Before freezing, vesicles and each solute were distributed uniformly, while the ice formation generated spatially heterogeneous patterns for both vesicles and solutes (Fig. S9). Pixel-wise analysis revealed strong positive correlations between rhodamine and calcein, GFP, or DNA fluorescence in the frozen state, indicating that membrane material and solutes were co-concentrated within the same confined regions (Fig. 3c). When we mapped fluorescence-rich pixels accounting for the upper 5% of the total fluorescence signal onto the corresponding grayscale bright-field images, fluorescence-rich regions substantially overlapped with the dark regions of the bright-field images, which we interpret as interstitial liquid regions between ice crystals (Figs. 3d and S10).

After thawing, GFP- and DNA-rich regions remained partially co-localized with rhodamine-rich micron-sized structures, suggesting that concentrated biomolecules were retained within or associated with membrane-rich compartments. In contrast, calcein fluorescence became more uniformly distributed and showed little apparent co-localization with rhodamine-rich structures, consistent with the lack of enrichment for calcein in GVs observed by confocal microscopy in Fig. 2 (Fig. S9). These observations suggest that freeze-induced molecular concentration does not necessarily translate into equivalent enrichment within GVs. Such differences may reflect molecule-dependent diffusion and interactions with membranes or other solutes during thawing and vesicle formation, potentially affecting molecular capture and retention within GVs.

### Slow FT assembles a functional transcription-translation system within vesicles

Having shown that multiple types of molecules can be co-concentrated with lipids during slow FT and incorporated into GVs, we next asked whether slow FT process could assemble a molecular system inside vesicles in its functional form. To test this, we used the PURE system, a reconstituted transcription–translation system, consisting of around 70 defined components, including ribosomes, enzymes, tRNAs, amino acids, and energy-regeneration components. Because its activity requires the coordinated presence of many molecular species with different sizes and physicochemical properties, the PURE system provides a stringent test of whether slow FT can assemble complex biochemical functionality within membrane compartments.

100 nm empty vesicles were mixed with the undiluted PURE system and DNA encoding GFP, and then subjected to slow FT. After slow FT, RNase was added to suppress gene expression in the extravesicular solution. The samples were then incubated at 37 °C for 4 h, and GFP fluorescence was measured as a readout of protein synthesis inside intact vesicles (Fig. 4a).

**Figure 4.**
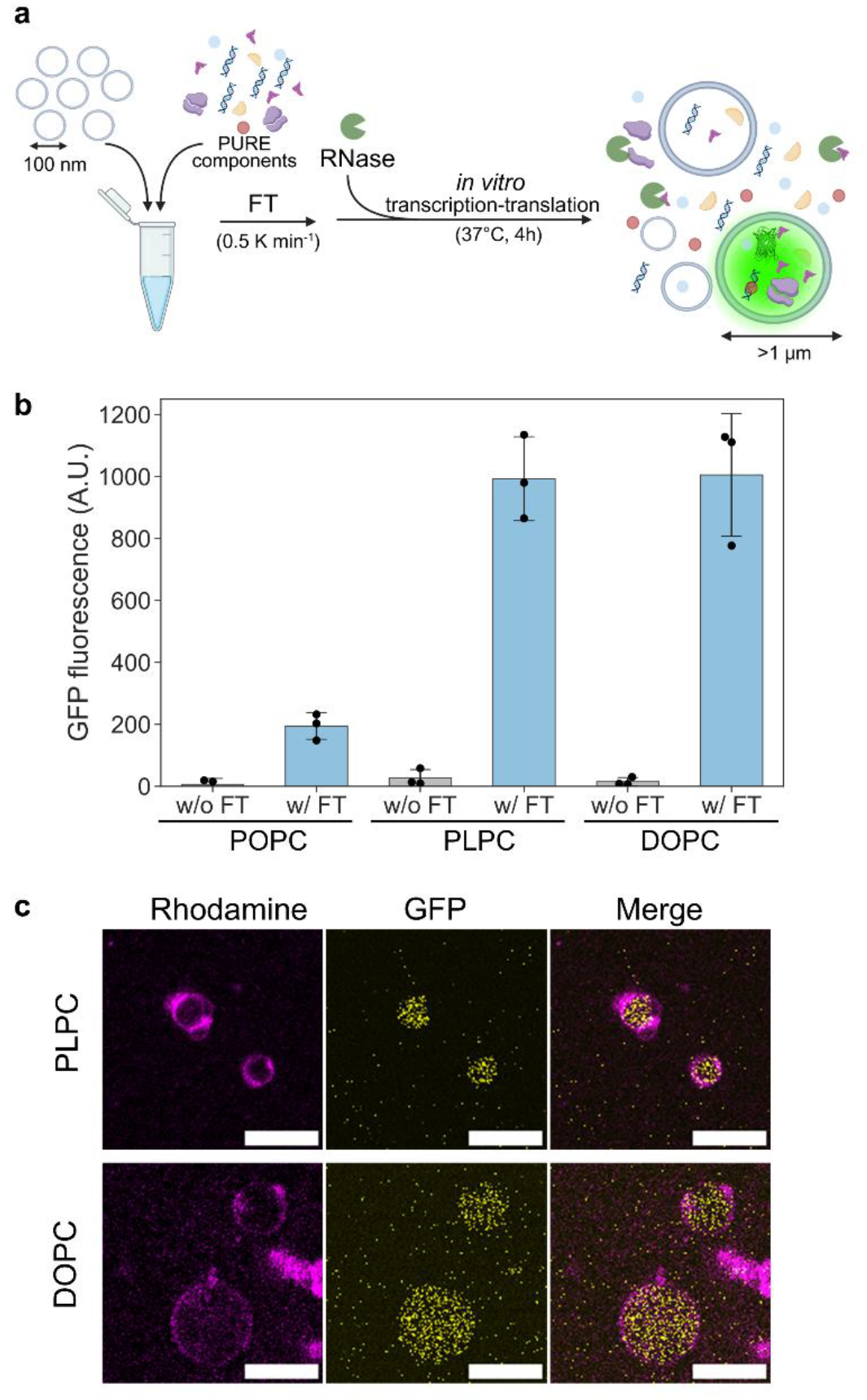
Slow FT assembles a functional transcription-translation system (PURE system) within vesicles. (a) Schematic illustration of FT-induced assembly of PURE system inside vesicles. In contrast to the preceding experiments using individual model solutes in HEPES buffer, vesicles were subjected to FT in the presence of the multicomponent PURE system, including the DNA template. (b) GFP fluorescence intensity in POPC, PLPC and DOPC vesicles with and without slow FT. Data points represent independent experiments (*n=*3), and error bars indicate mean ± s.d. (c) Representative confocal fluorescence images of PLPC and DOPC vesicles after slow FT, showing GFP expression inside GVs. Scale bar: 10 µm.

GFP expression was detected after slow FT for all three lipid compositions (POPC, PLPC and DOPC), but was more pronounced in PLPC and DOPC vesicles than in POPC vesicles (Fig. 4b). Confocal imaging showed GFP fluorescence within GVs, indicating that GV formation was accompanied by the co-encapsulation of multiple PURE components, including large macromolecular assemblies such as ribosomes (Fig. 4c). Thus, slow FT can encapsulate not only individual biomolecules but also a multicomponent biochemical system within GVs in a functional form, while physically protecting the encapsulated reaction from externally added RNase.

### Slow FT activates a diluted PURE system within vesicles

Having shown that slow FT can enrich diverse molecular species and encapsulate a functional multicomponent PURE system within vesicles, we finally explored whether slow FT could activate a diluted PURE system that was poorly active in a bulk solution by concentrating its components. Empty DOPC vesicles were mixed with 1×, 0.4× or 0.3× PURE reaction mixtures containing GFP-encoding DNA (undiluted, 2.5-fold diluted and 3.33-fold diluted with milli-Q water, respectively), and subjected to slow FT (Fig. 5a). Bulk measurements of GFP production across a dilution series of the PURE system showed that GFP production at 0.4× and 0.3× PURE fell to, or below, the detection limit (Fig. 5b, dashed line). Strikingly, after slow FT, confocal microscopy imaging revealed GFP production selectively within GVs at both 0.4× and 0.3× PURE, whereas little detectable expression was observed in the surrounding solution (Fig. 5c). Unlike in Fig. 4, the external solution was not treated with RNase; thus, GFP synthesis could in principle have occurred outside the vesicles had the diluted PURE system remained active in bulk. However, the selective GFP production inside the vesicles indicates that activity was restored specifically within the compartments. Giant vesicles prepared by the emulsion-transfer method^42,43^, for which molecular enrichment is not expected, showed no detectable GFP fluorescence either inside the vesicles or in the surrounding solution at 0.3× PURE (Fig. S11). Compartmentalization alone was therefore insufficient to activate the diluted reaction, indicating that slow FT-mediated molecular enrichment was critical for restoring gene expression.

**Figure 5.**
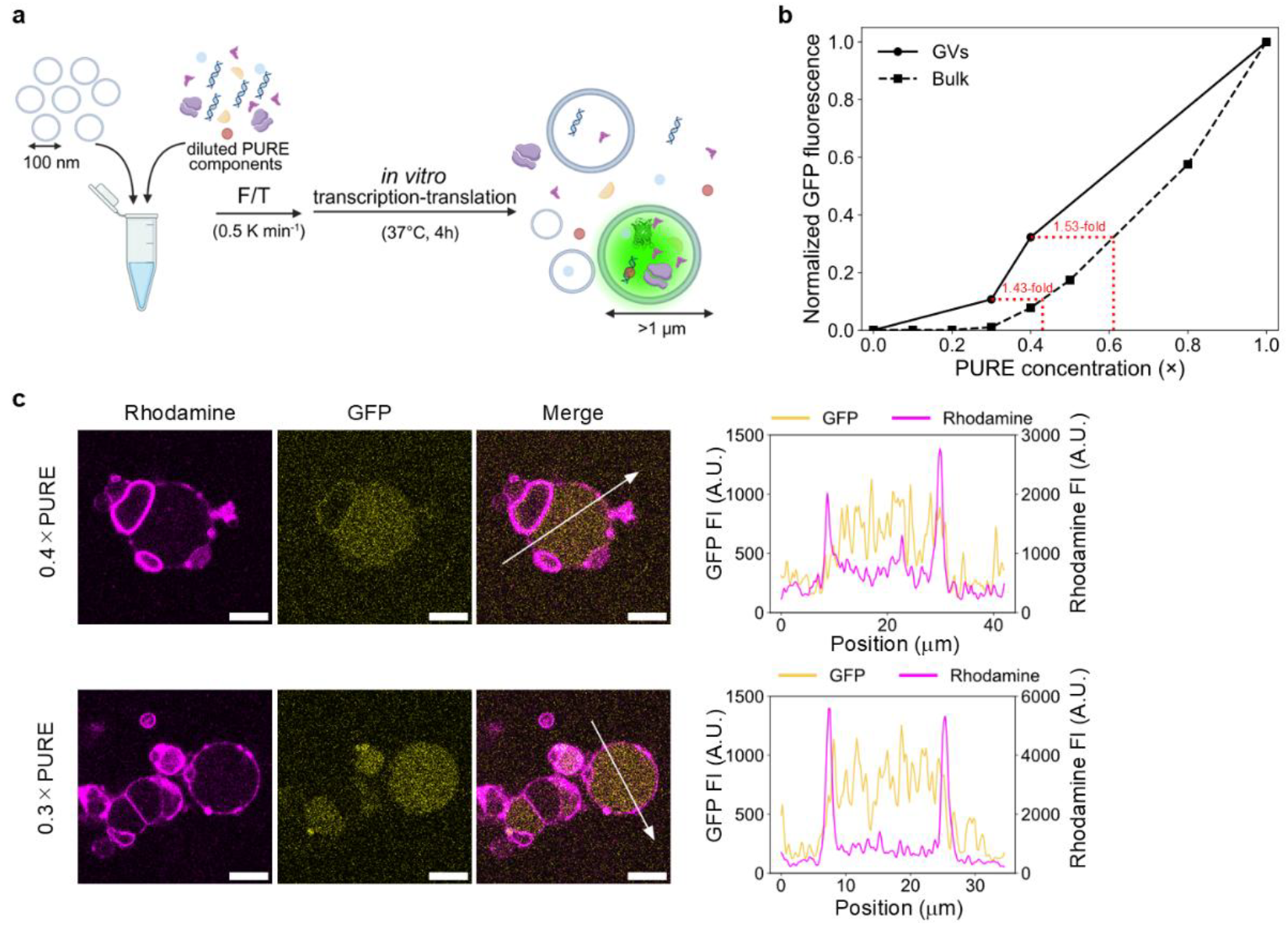
Slow FT activates a diluted PURE system within GVs. (a) Schematic illustration of experimental design. (b) GFP output as a function of PURE concentration in bulk reactions (dashed line) and FT-generated GVs (solid line), with each dataset normalized independently to its respective 1× PURE condition. For the GV system, 10,000 events were analyzed by flow cytometry, and the total GFP fluorescence summed across all events was normalized to the 1× PURE condition and plotted. See Fig. S12 for the gating strategy and detailed flow cytometry analysis. (c) Representative confocal fluorescence images and corresponding line intensity profiles of vesicles after slow FT from 0.4× and 0.3× PURE mixtures. Images used for line-profile analysis were processed with a Gaussian blur of σ = 1 pixel to reduce noise. Scale bar: 10 µm.

Flow cytometry further confirmed the reactivation of the diluted PURE system within FT-generated GVs. Among 10,000 analyzed GV events, GVs exhibiting GFP fluorescence (GFP-positive GVs) accounted for 63.2%, 32.6% and 19.7% of the populations generated with 1×, 0.4× and 0.3× PURE, respectively (Fig. S12). To quantify GFP production at the population level, we calculated the total GFP output from the fraction of GFP-positive GVs, and normalized the resulting values to the 1× PURE condition. Plotting this output against the initial PURE concentration yielded the dose–response relationship for the GV system (Fig. 5b, solid line). GFP production in FT-generated GVs declined more gradually upon dilution than in bulk solution. In particular, the activities observed with 0.4× and 0.3× PURE in GVs corresponded to those expected for approximately 0.61× (1.53-fold relative to 0.4×) and 0.43× (1.43-fold relative to 0.3×) PURE in bulk, respectively, corresponding to an effective concentration shift of approximately 1.5-fold (Fig. 5b). In conclusion, our results demonstrate that an approximately 1.5-fold concerted enrichment of diverse molecular components can shift the local reaction environment across a functional threshold and, when coupled to encapsulation, transform an otherwise inactive molecular system into functional protocellular compartments.

## Conclusion

The emergence of cellular life occurred during a chapter of Earth’s history that the geological record has largely erased^44^. We have therefore sought to reconstruct this lost chapter through bottom-up approaches, asking from both environmental and prebiotic-chemical perspectives what such environments were physically capable of doing.

A wide range of prebiotic chemistry leading to the formation of life’s molecular building blocks has been reported under markedly different conditions, including atmospheric reactions^45^, hydrothermal systems^46^, and freezing environments^47^. This diversity raises the possibility that cellular life did not emerge within a single setting, but through repeated dispersal and reconcentration of molecules produced across different environments^5^.

Among plausible prebiotic settings capable of concentrating dispersed molecules^10–21,28,29^, our results highlight FT as a physical process capable of assembling complex multicomponent systems and directly converting them into functional, cell-sized membrane compartments. Previous studies have successfully demonstrated prebiotically relevant chemical processes under FT conditions such as nucleobase synthesis^22^, nonenzymatic template-directed RNA copying^26^, ribozyme replication^23^, functional ribozyme assembly^25^, and DNA hybridization^27^. The PURE system used here, by contrast, comprises approximately 70 defined components operating through multiple coupled reaction steps, extending FT-driven chemistry to a substantially greater level of molecular and reaction-network complexity. Moreover, unlike approaches that induce biochemical function through content mixing between preformed GUVs^48–50^, FT here couples compartment formation and concentration-driven biochemical activation within a single physical process, suggesting a route to the direct formation of functional compartments without requiring preformed cellular compartments.

Molecular encapsulation by slow FT was also compositionally selective. Although chemically distinct solutes were co-concentrated within ice interstices, their final enrichment within GVs differed markedly: proteins and DNA were retained more efficiently than calcein (Fig. 2). Slow FT also generated substantial vesicle-to-vesicle heterogeneity. Long-tailed enrichment distributions (Fig. 2) and rare, exceptionally GFP-rich GVs (Fig. S13) revealed broad variation in molecular composition and functional output among individual compartments. Compartmentalization may therefore convert microscopic fluctuations in molecular enrichment into discrete compositional and functional diversity. If such differences affect the persistence, growth, or reproduction of compartments over repeated environmental cycles, successive FT events could in principle expose this diversity to selection, linking environmental fluctuations to early protocellular evolution.

Overall, our findings establish temperature oscillations in icy environments as a physical pathway by which dilute and initially dispersed molecular components can be concentrated and organized into cell-sized functional compartments. As space exploration now turns towards the icy ocean worlds of our Solar System^51,52^, these findings invite a broader view of ice—not merely as a passive backdrop or barrier to life, but as a dynamic physical medium capable of organizing matter towards cell-like organization and function.

## Materials and methods

### Chemicals

1-palmitoyl-2-oleoyl-glycero-3-phosphocholine (POPC) and 1,2-dioleoyl-sn-glycero-3-phosphocholine (DOPC) were purchased from NOF Corporation. 1-palmitoyl-2-linoleoyl-sn-glycero-3-phosphocholine (PLPC) and 1,2-dioleoyl-sn-glycero-3-phosphoethanolamine-N-(lissamine rhodamine B sulfonyl)(ammonium salt) (18:1 Liss Rhod PE) were purchased from Avanti Polar Lipids. Calcein and cobalt(II) chloride hexahydrate were purchased from FUJIFILM Wako Pure Chemical Corporation. Ovalbumin Alexa Fluor 647 conjugate (OA647) and SYBR Gold nucleic acid stain were purchased from Thermo Fisher Scientific. 5’-FITC labelled single-stranded DNA oligonucleotides consisting of AC-repeat sequences with lengths of 10, 20, 40 and 80 nucleotides were purchased from FASMAC. PURE*frex***®**1.0 was obtained from GeneFrontier Corporation. Ribonuclease A (RNase A) and HEPES were purchased from Sigma-Aldrich. SNAP-tagged GFP was purified as previously reported^53^. A plasmid encoding GFP was constructed by amplifying the green fluorescent protein gene^54^ from pUC-frGFP (Musaiboukun; Taiyo Nippon Sanso) and assembling it into the pIVEX2.3 vector using the NEBuilder HiFi DNA Assembly Kit (New England Biolabs). The plasmid sequence was verified by Sanger sequencing. DNA templates encoding GFP used in this study were PCR products amplified from this plasmid using primers T7F (5’-ATGCGTCCGGCGTAGAGGATCGAGA-3’) and T7R (5’-AGGGGTTATGCTAGTTATTGCTCAGCGG-3’), and KOD polymerase (Toyobo) according to the manufacturer’s instructions.

### Preparation of empty vesicles with an average diameter of 100 nm

~100 nm size large unilamellar vesicles (LUVs) were prepared by extrusion methodology^55^ with some modification. Briefly, 10 mg of lipid dissolved in ~200 µl of pure chloroform was rotated and evaporated under reduced pressure in a round-bottom flask for 1 h. For fluorescence visualization, 0.1 mol% 18:1 Liss Rhod PE was included in the lipid mixture before film formation. The resulting dried lipid film was hydrated with 300 µl of 50 mM HEPES buffer (pH 7.4), to yield 300 µl of ~30 mg/mL vesicle solution (multilamellar vesicles). The vesicle solution was sonicated (VS-1003, AS ONE Corporation) for 10 min (repeated for a total of 10 min: 28 kHz for 1min, 45 kHz for 1min, 100 kHz for 3s) and vortexed for 15 s to resuspend completely. The suspension was then subjected to 5 cycles of repeated FT (incubated in liquid nitrogen for 1 min, then incubated in 65°C of water bath until thawed). The vesicle suspension was extruded 21 times using an extruder (Avanti Polar Lipids) through a 100 nm polycarbonate membrane filter. Subsequently, centrifugal separation at 20,000 g for 25 min was conducted to eliminate aggregates. The phospholipid concentration was determined using LabAssay^™^ Phospholipid kit (Choline Oxidase DAOS method, FUJIFILM Wako Pure Chemical Corporation) according to manufacturers’ instruction. LUVs suspensions were diluted to the indicated lipid concentrations before use.

### Freeze-thaw (FT) treatment

Empty vesicles were mixed with the indicated solutes in 50 mM HEPES buffer (pH 7.4) in 0.2-ml PCR tubes to a final volume of 20 µl. Unless otherwise stated, the final lipid concentration was 1 mg ml^-1^. The final solute concentrations were 10 µM calcein, 1 µM SNAP-GFP, 1 µM OA647, 10 nM DNA (GPF encoding DNA template) with 0.2× SYBR Gold, or 1 µM 5’-FITC labelled AC-repeat ssDNA oligonucleotides of 10, 20, 40 or 80 nucleotides. In reactions containing the PURE system, empty vesicles were mixed with PURE*frex*1.0 and 10 nM GFP-encoding DNA. PURE components mixture and DNA were both diluted with Milli-Q water when required.

The prepared samples were subjected to a single freeze–thaw cycle under the conditions listed in Table 1. For fast freezing, samples were immersed in liquid nitrogen for 1 min. For moderate freezing, samples were incubated for 10 min on a metal block pre-cooled in a −30 °C freezer. For fast thawing, samples were incubated on a metal incubator set to 60 °C. For moderate thawing, samples were incubated on a metal block placed on ice. Slow freezing and slow thawing were performed using a programmable circulating low-temperature bath (PCC-7000; EYELA, Tokyo Rikakikai), with cooling from 4 °C to −27.5 °C and warming from −27.5 °C to 4 °C at 0.5 K min^-1^.

For calcein encapsulation assays, external calcein was quenched after FT by adding 40 µl of 100 µM CoCl_2_ solution to 20 µl of sample, and the remaining membrane-protected calcein fluorescence was measured using a microplate reader (BioTek Synergy H1, Agilent Technologies). A vesicle sample without FT treatment was used as a negative control to estimate the background calcein fluorescence remaining after CoCl_2_ addition. Calcein encapsulation (%) was calculated as:

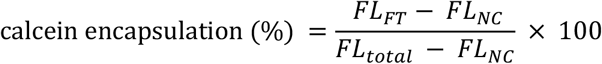

where *FL*_*FT*_ is the calcein fluorescence intensity of the FT treated sample after CoCl_2_ addition, *FL*_*NC*_ is the calcein fluorescence intensity of the sample without FT treatment after CoCl_2_ addition (negative control), and *FL*_*total*_is the calcein fluorescence intensity corresponding to the total amount of calcein initially added to the sample.

The theoretical calcein encapsulation value for uniformly dispersed 100 nm vesicles was calculated as the ratio of the total internal aqueous volume of input vesicle population to the sample volume. Vesicles were assumed to be spherical bilayer vesicles with an outer diameter of 100 nm, a membrane thickness of 3.9 nm and an area per lipid molecule of 0.64 nm^2^, with a lipid molecular weight of 760 g mol^-156^. For a 20 µl sample containing 1 mg ml^-1^ lipid, this calculation gave a theoretical encapsulation value of 0.36%.

For PURE system experiments, samples were incubated at 37 °C for 4 h after FT. Where indicated, 1 µl of RNase A solution (5 mg ml^-1^) was added to suppress gene expression in the external solution.

### Dynamic light scattering analysis (DLS)

The size distribution of vesicles was determined by dynamic light scattering analysis (DelsaMax CORE, Beckman Coulter). Measurements were conducted using a laser wavelength of 660.9 nm, and the temperature in the instrument was kept at 25°C. For each sample, three measurements of 10 runs were performed, and the histogram was generated using the average DLS intensity across these three measurements.

### Bulk fluorescence measurements

Bulk fluorescence measurements were performed using a microplate reader (BioTek Synergy H1, Agilent Technologies). Calcein fluorescence and GFP fluorescence expressed by the PURE system were measured with excitation at 490 nm and emission at 520 nm.

### *In situ* observations

*In situ* bright-field and fluorescence imaging during FT was performed using an inverted fluorescence microscope (IX73, Olympus) equipped with a Peltier cooling/heating stage (Linkam, model 10021). Samples were placed on a glass slide with a spacer, covered with a coverslip, mounted on the cooling stage and subjected to the temperature profile described in the main text. Bright-field and fluorescence time-lapse images were acquired during the FT process. Fluorescence images of calcein, GFP and SYBR Gold-stained DNA were acquired using a BP460–495 excitation filter, DM505 dichroic mirror and BA510IF emission filter. Rhodamine fluorescence was acquired using a BP530–550 excitation filter, DM570 dichroic mirror and BA575IF emission filter. To minimize frost formation during cooling, the cooling stage was placed within a transparent plastic enclosure that was continuously purged with N_2_ gas.

### Confocal laser scanning microscopy

Confocal images of vesicles were acquired using a confocal laser scanning microscope LSM 900 (Carl Zeiss), equipped with a 63×/1.40 oil immersion objective at room temperature. The fluorescence imaging was performed with excitation wavelength of 488 nm for calcein, GFP, SYBR gold-stained DNA, FITC-labelled ssDNA, 561 nm for Rhodamine, and 640 nm for OA647. For quantitative comparisons, images within the same experiment were acquired under identical laser power, detector gain and imaging settings. Images were analyzed using Fiji/ImageJ. Contrast and brightness were adjusted only for visualization, and all quantitative analyses were performed using raw image data.

### Flowcytometric analysis

Vesicle populations were analyzed by flow cytometry using a FACSAria III cell sorter (BD Biosciences). Samples were diluted before measurement in 50 mM HEPES buffer (pH 7.4). For samples containing PURE system, dilution was performed using an osmotically matched buffer containing HEPES, potassium glutamate and magnesium acetate. A total of 10,000 events were collected for each sample unless otherwise indicated. Calcein and GFP fluorescence were detected using a 488-nm blue laser and a 530/30-nm emission filter, whereas rhodamine fluorescence was detected using a 561-nm yellow-green laser and a 585/15-nm emission filter. Detector voltages and thresholds were optimized for each experiment and kept constant within the same experiment. Detailed voltage and threshold settings and gating strategies are provided in the Supplementary Information. Data were analyzed using FlowJo software (v10.10.0; BD Biosciences).

## Data availability

The data supporting the findings of this study are included in the Article and its Supplementary Information. Additional information is available from the corresponding author upon request.

## Acknowledgments

This study was supported by Human Frontier Science Program Grant Number RGP003/2023 (T.M.), JSPS KAKENHI Grant Numbers 22K21344 (T.M.), 21H05228 (T.M.), JST ASPIRE Grant Number JPMJAP24B4 (T.M.) and JST SPRING Grant Number JPMJSP JPMJSP2180 (TS). We thank Liam M. Longo (ELSI, Institute of Science Tokyo) for critically reading the manuscript and providing constructive comments.

## Author Contributions

T.S. performed most of the experiments. K.P.S. contributed to the experimental work during the early phase of the project. T.S., S.H., H.-E.L., R.N. and T.M. performed and contributed to the design and interpretation of the *in situ* observation experiments. T.S., K.P.S., N.N. and T.M. conceived the project. T.S. and T.M. wrote the manuscript, with contributions from all authors. All authors discussed the results and reviewed and edited the manuscript.

## Notes

Competing interests

The authors declare no competing interests.

## References

1. Szostak, J. W., Bartel, D. P. & Luisi, P. L. Synthesizing life. Nature 409, 387–390 (2001).

2. Ellis, R. J. Macromolecular crowding: obvious but underappreciated. Trends Biochem. Sci. 26, 597–604 (2001).

3. Patel, B. H., Percivalle, C., Ritson, D. J., Duffy, C. D. & Sutherland, J. D. Common origins of RNA, protein and lipid precursors in a cyanosulfidic protometabolism. Nat. Chem. 7, 301–307 (2015).

4. Powner, M. W., Gerland, B. & Sutherland, J. D. Synthesis of activated pyrimidine ribonucleotides in prebiotically plausible conditions. Nature 459, 239–242 (2009).

5. Kitadai, N. & Maruyama, S. Origins of building blocks of life: A review. Geosci. Front. 9, 1117–1153 (2018).

6. Cleaves, H. J. Prebiotic chemistry: geochemical context and reaction screening. Life (Basel) 3, 331–345 (2013).

7. Baaske, P. et al. Extreme accumulation of nucleotides in simulated hydrothermal pore systems. Proc. Natl. Acad. Sci. U. S. A. 104, 9346–9351 (2007).

8. Sarkar, S. et al. Prebiological membranes and their role in the emergence of early cellular life. J. Membr. Biol. 253, 589–608 (2020).

9. Mizuuchi, R. & Ichihashi, N. Primitive compartmentalization for the sustainable replication of genetic molecules. Life (Basel) 11, 191 (2021).

10. Rajamani, S. et al. Lipid-assisted synthesis of RNA-like polymers from mononucleotides. Orig. Life Evol. Biosph. 38, 57–74 (2008).

11. Mamajanov, I. et al. Ester formation and hydrolysis during wet–dry cycles: Generation of far-from-equilibrium polymers in a model prebiotic reaction. Macromolecules 47, 1334–1343 (2014).

12. Da Silva, L.Maurel, M.-C. & Deamer, D. Salt-promoted synthesis of RNA-like molecules in simulated hydrothermal conditions. J. Mol. Evol. 80, 86–97 (2015).

13. Becker, S. et al. Wet-dry cycles enable the parallel origin of canonical and non-canonical nucleosides by continuous synthesis. Nat. Commun. 9, 163 (2018).

14. Song, X., Simonis, P., Deamer, D. & Zare, R. N. Wet-dry cycles cause nucleic acid monomers to polymerize into long chains. Proc. Natl. Acad. Sci. U. S. A. 121, e2412784121 (2024).

15. Ianeselli, A. et al. Water cycles in a Hadean CO2 atmosphere drive the evolution of long DNA. Nat. Phys. 18, 579–585 (2022).

16. Salditt, A. et al. Ribozyme-mediated RNA synthesis and replication in a model Hadean microenvironment. Nat. Commun. 14, 1495 (2023).

17. Dirscherl, C. F. et al. A heated rock crack captures and polymerizes primordial DNA and RNA. Phys. Chem. Chem. Phys. 25, 3375–3386 (2023).

18. Dass, A. V. et al. RNA oligomerisation without added catalyst from 2’,3’-cyclic nucleotides by drying at air-water interfaces. ChemSystemsChem 5, (2023).

19. Matreux, T., Aikkila, P., Scheu, B., Braun, D. & Mast, C. B. Heat flows enrich prebiotic building blocks and enhance their reactivity. Nature 628, 110–116 (2024).

20. Schwintek, P., Eren, E., Mast, C. B. & Braun, D. Prebiotic gas flow environment enables isothermal nucleic acid replication. Elife 13, (2025).

21. Floroni, A. et al. Membraneless protocell confined by a heat flow. Nat. Phys. 21, 1303–1310 (2025).

22. Kanavarioti, A., Monnard, P. A. & Deamer, D. W. Eutectic phases in ice facilitate nonenzymatic nucleic acid synthesis. Astrobiology 1, 271–281 (2001).

23. Attwater, J., Wochner, A., Pinheiro, V. B., Coulson, A. & Holliger, P. Ice as a protocellular medium for RNA replication. Nat. Commun. 1, 76 (2010).

24. Menor-Salván, C. & Marín-Yaseli, M. R. Prebiotic chemistry in eutectic solutions at the water-ice matrix. Chem. Soc. Rev. 41, 5404–5415 (2012).

25. Mutschler, H., Wochner, A. & Holliger, P. Freeze-thaw cycles as drivers of complex ribozyme assembly. Nat. Chem. 7, 502–508 (2015).

26. Zhang, S. J., Duzdevich, D., Ding, D. & Szostak, J. W. Freeze-thaw cycles enable a prebiotically plausible and continuous pathway from nucleotide activation to nonenzymatic RNA copying. Proc. Natl. Acad. Sci. U. S. A. 119, e2116429119 (2022).

27. Noda, N. et al. Slow freeze - thaw cycles enhanced hybridization of kilobase DNA with long complementary sticky ends. ChemSystemsChem 6, (2024).

28. Ferris, J. P., Hill, A. R., Jr, Liu, R. & Orgel, L. E. Synthesis of long prebiotic oligomers on mineral surfaces. Nature 381, 59–61 (1996).

29. Hazen, R. M. & Sverjensky, D. A. Mineral surfaces, geochemical complexities, and the origins of life. Cold Spring Harb. Perspect. Biol. 2, a002162 (2010).

30. Zahnle, K. et al. Emergence of a habitable planet. Space Sci. Rev. 129, 35–78 (2007).

31. Kadoya, S., Krissansen-Totton, J. & Catling, D. C. Probable cold and alkaline surface environment of the Hadean Earth caused by impact ejecta weathering. Geochem. Geophys. Geosyst. 21, (2020).

32. Fox-Powell, M. G. & Cousins, C. R. Partitioning of crystalline and amorphous phases during freezing of simulated Enceladus ocean fluids. J. Geophys. Res. Planets 126, e2020JE006628 (2021).

33. Cleaves, H. J., 2nd et al. Mineral-organic interfacial processes: potential roles in the origins of life. Chem. Soc. Rev. 41, 5502–5525 (2012).

34. Shimizu, Y. et al. Cell-free translation reconstituted with purified components. Nat. Biotechnol. 19, 751–755 (2001).

35. Shimizu, Y., Kanamori, T. & Ueda, T. Protein synthesis by pure translation systems. Methods 36, 299–304 (2005).

36. Chiba, M., Miyazaki, M. & Ishiwata, S. Quantitative analysis of the lamellarity of giant liposomes prepared by the inverted emulsion method. Biophys. J. 107, 346–354 (2014).

37. Shinoda, T. et al. Compositional selection of phospholipid compartments in icy environments drives the enrichment of encapsulated genetic information. Chem. Sci. 16, 23321–23329 (2025).

38. Poojari, C. S., Scherer, K. C. & Hub, J. S. Free energies of membrane stalk formation from a lipidomics perspective. Nat. Commun. 12, 6594 (2021).

39. Kumarage, T. et al. Cholesterol modulates membrane elasticity via unified biophysical laws. Nat. Commun. 16, 7024 (2025).

40. Blicher, A., Wodzinska, K., Fidorra, M., Winterhalter, M. & Heimburg, T. The temperature dependence of lipid membrane permeability, its quantized nature, and the influence of anesthetics. Biophys. J. 96, 4581–4591 (2009).

41. Peter, B. & Schwille, P. Elucidating the mechanism of freeze-thaw driven content mixing between protocells. ChemSystemsChem http://doi.org/10.1002/syst.202300008 (2023) doi:10.1002/syst.202300008.

42. Pautot, S., Frisken, B. J. & Weitz, D. A. Production of unilamellar vesicles using an inverted emulsion. Langmuir 19, 2870–2879 (2003).

43. Nishimura, K. et al. Cell-free protein synthesis inside giant unilamellar vesicles analyzed by flow cytometry. Langmuir 28, 8426–8432 (2012).

44. Westall, F., Brack, A., Fairén, A. G. & Schulte, M. D. Setting the geological scene for the origin of life and continuing open questions about its emergence. Front. Astron. Space Sci. 9, 1095701 (2023).

45. Miller, S. L. A production of amino acids under possible primitive earth conditions. Science 117, 528–529 (1953).

46. Huber, C. & Wächtershäuser, G. alpha-Hydroxy and alpha-amino acids under possible Hadean, volcanic origin-of-life conditions. Science 314, 630–632 (2006).

47. Schwartz, A. W., Joosten, H. & Voet, A. B. Prebiotic adenine synthesis via HCN oligomerization in ice. Biosystems. 15, 191–193 (1982).

48. Tsuji, G., Fujii, S., Sunami, T. & Yomo, T. Sustainable proliferation of liposomes compatible with inner RNA replication. Proc. Natl. Acad. Sci. U. S. A. 113, 590–595 (2016).

49. Salibi, E., Peter, B., Schwille, P. & Mutschler, H. Periodic temperature changes drive the proliferation of self-replicating RNAs in vesicle populations. Nat. Commun. 14, 1222 (2023).

50. Abil, Z. et al. Darwinian evolution of self-replicating DNA in a synthetic protocell. Nat. Commun. 15, 9091 (2024).

51. Pappalardo, R. T. et al. Science overview of the Europa Clipper mission. Space Sci. Rev. 220, (2024).

52. Grasset, O. et al. JUpiter ICy moons Explorer (JUICE): An ESA mission to orbit Ganymede and to characterise the Jupiter system. Planet. Space Sci. 78, 1–21 (2013).

53. Uyeda, A., Watanabe, T., Hohsaka, T. & Matsuura, T. Different protein localizations on the inner and outer leaflet of cell-sized liposomes using cell-free protein synthesis. Synth. Biol. 3, ysy007 (2018).

54. Waldo, G. S., Standish, B. M., Berendzen, J. & Terwilliger, T. C. Rapid protein-folding assay using green fluorescent protein. Nat. Biotechnol. 17, 691–695 (1999).

55. Mayer, L. D., Hope, M. J. & Cullis, P. R. Vesicles of variable sizes produced by a rapid extrusion procedure. Biochim. Biophys. Acta 858, 161–168 (1986).

56. Kučerka, N.Nieh, M.-P. & Katsaras, J. Fluid phase lipid areas and bilayer thicknesses of commonly used phosphatidylcholines as a function of temperature. Biochim. Biophys. Acta 1808, 2761–2771 (2011).

