## Supplementary Information for "Ice drives the emergence of functional protocells"

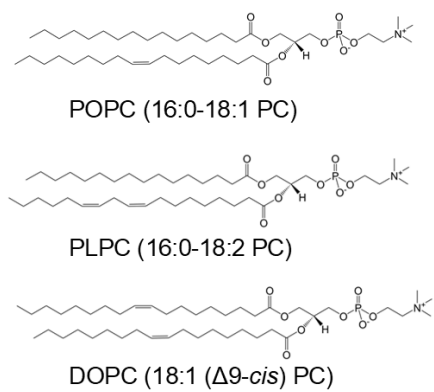

**Figure S1. Chemical structures of the phospholipids used in this study.** 1-palmitoyl-2-oleoyl-glycero-3-phosphocholine (POPC, 16:0-18:1 PC), 1-palmitoyl-2-linoleoyl-sn-glycero-3-phosphocholine (PLPC, 16:0-18:2 PC), and 1,2-dioleoyl-sn-glycero-3-phosphocholine (DOPC, 18:1 ($\Delta^9$-Cis) PC).

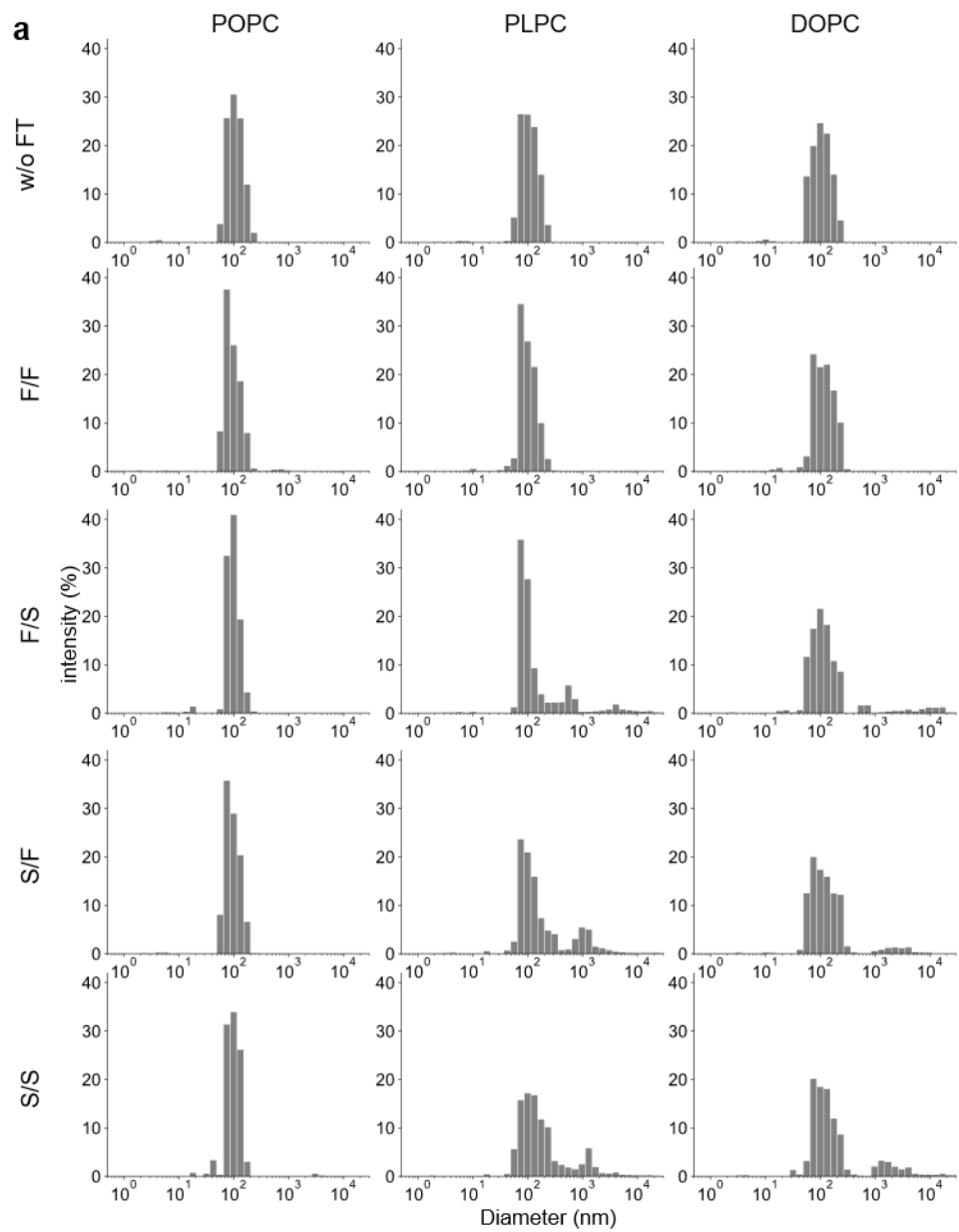

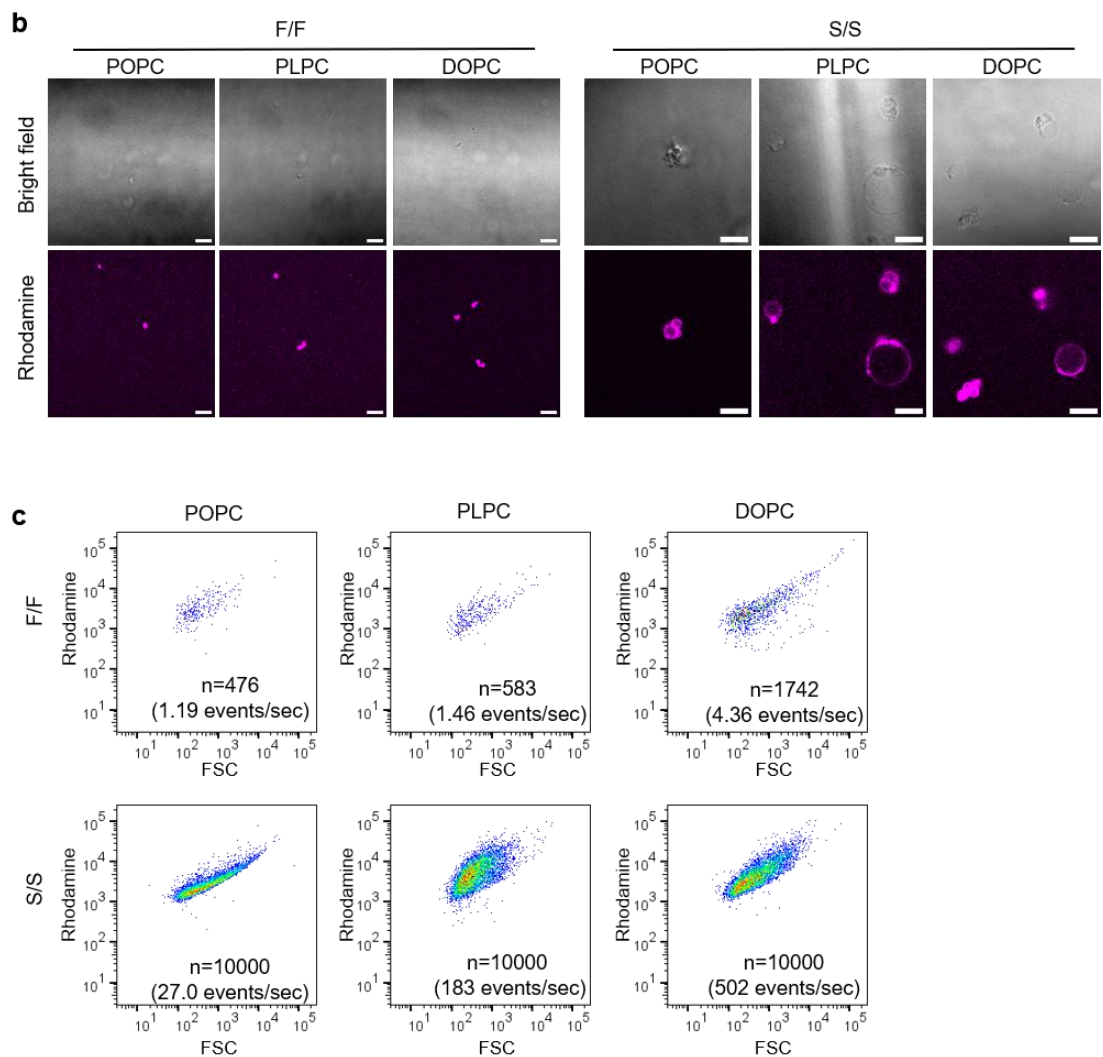

**Figure S2. Effects of freeze-thaw (FT) conditions on giant vesicle (GV) formation.** (a) Rhodamine-labelled empty 100 nm vesicles composed of POPC, PLPC or DOPC were subjected to different FT conditions, including fast freezing/fast thawing (F/F), fast freezing/slow thawing (F/S), slow freezing/fast thawing (S/F) and slow freezing/slow thawing (S/S), as defined in Table 1. Vesicle size distributions before and after FT treatment were analyzed by dynamic light scattering (DLS). Before FT, all vesicle samples showed a size distribution centered around 100 nm in diameter. After FT, PLPC and DOPC vesicles showed the appearance of micrometer-sized fractions, particularly under F/S, S/F and S/S conditions, whereas such enlarged fractions were not observed under F/F. In contrast, POPC vesicles showed little formation of enlarged fractions under the tested FT conditions. (b) Confocal microscopy was used to confirm the formation of GV's after FT. Under F/F conditions, only small aggregates or diffraction-limited fluorescent puncta were detectable. In contrast, S/S generated GV's for all lipid compositions, although POPC produced far fewer GV's than PLPC and DOPC. Scale bar: 10 μm. (c) The abundance of GV's was further quantified by flow cytometry. Detector voltages were set to 5, 350, 300 for forward scatter (FSC), side scatter (SSC), and rhodamine, respectively, with FSC and SSC thresholds both set to 200. Under these detection settings, flow cytometry mainly detected GV-sized fluorescent events, allowing the relative abundance of GV's to be compared by the event rate measured over a fixed acquisition time. Under F/F conditions, POPC, PLPC and DOPC showed event rates of 1.19, 1.46 and 4.36 events $s^{-1}$, respectively. In contrast, S/S markedly increased the event rates to 27.0, 183 and 502 events $s^{-1}$ for POPC, PLPC and DOPC, respectively. These results show that F/F produced few detectable GV's, whereas S/S strongly promoted GV formation. The extent of GV formation was also strongly dependent on lipid composition, with DOPC and PLPC producing substantially more GV's than POPC.

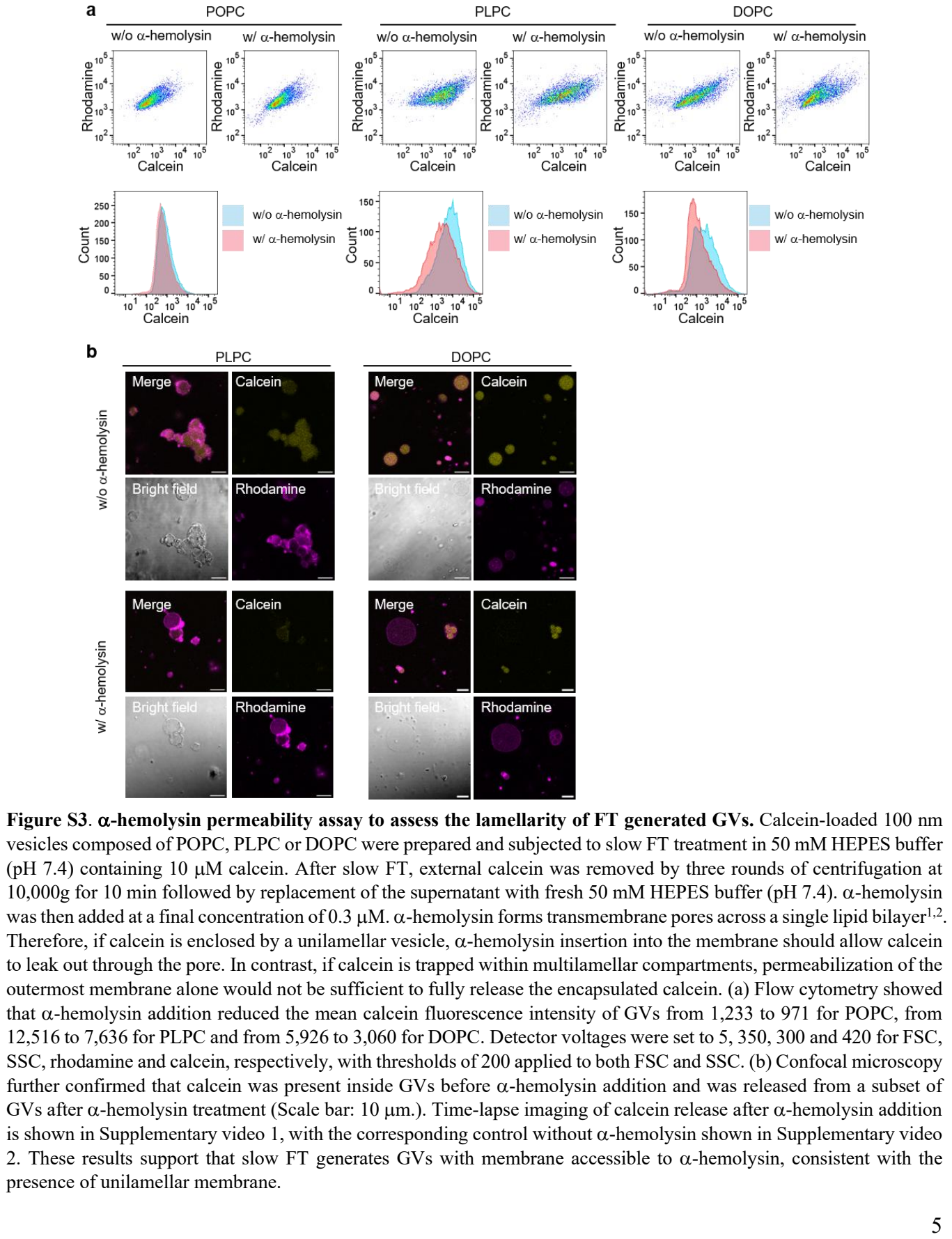

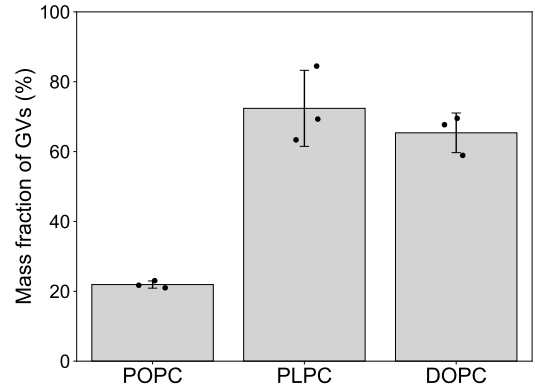

**Figure S4. Quantification of mass fraction of GVs after slow FT.** The mass fraction of GVs generated after slow FT was quantified for POPC, PLPC and DOPC vesicles. After slow FT, samples were centrifuged at $20,000 \times g$ for 25 min to separate GVs from non-enlarged vesicles; the initial 100 nm vesicles remained in the supernatant under this condition. The mass fraction of GVs was defined as the lipid mass recovered in the pellet fraction relative to the total lipid mass before centrifugation. The mass fraction of GVs were $21.9 \pm 1.03\%$, $72.4 \pm 10.9\%$ and $65.4 \pm 5.7\%$ for POPC, PLPC, and DOPC, respectively (mean ± s.d., $n = 3$). These results show that PLPC and DOPC vesicles grew to a similar extent under slow FT conditions, whereas POPC showed a substantially less growth.

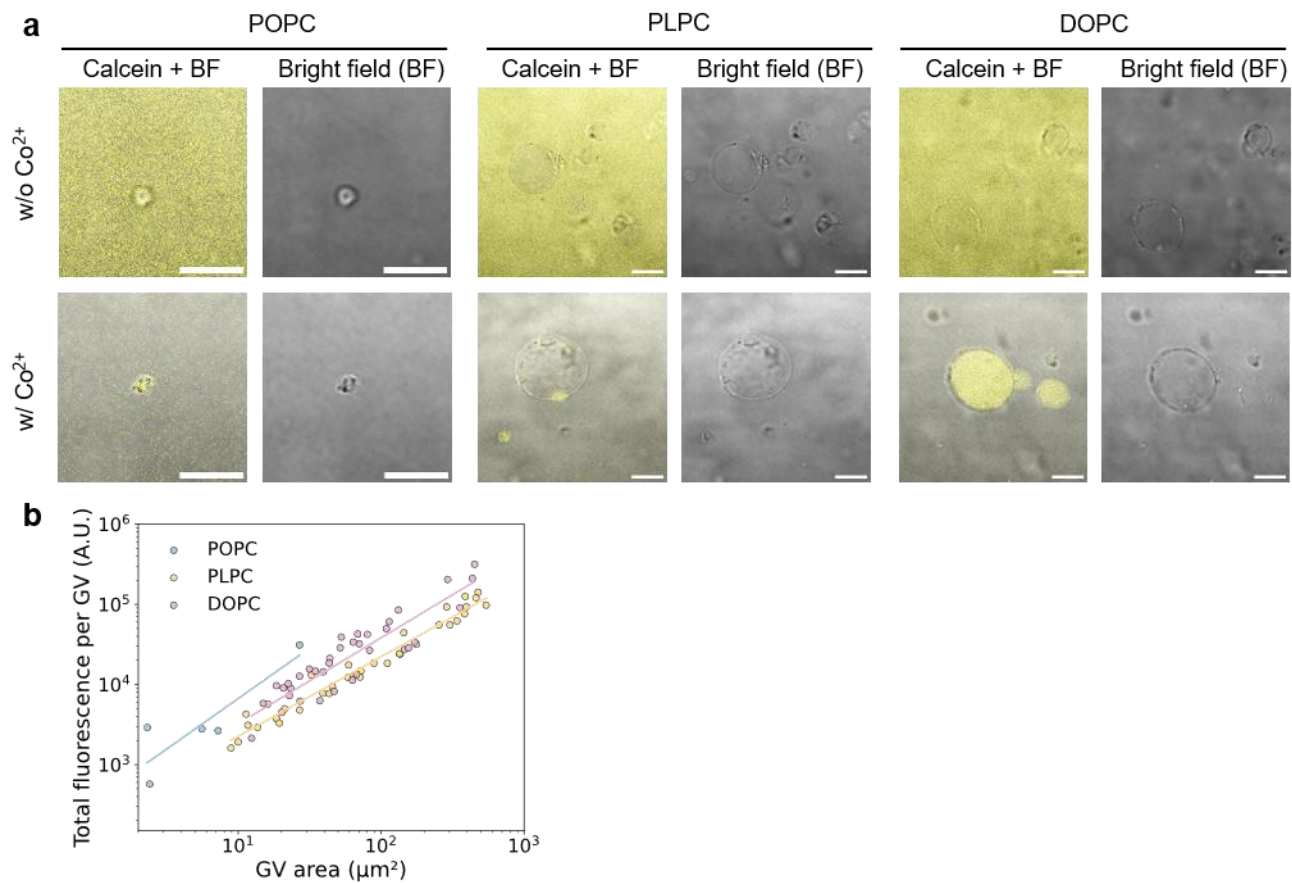

**Figure S5. Confocal imaging of calcein encapsulation after slow FT and cobalt quenching.** (a) Calcein encapsulation after slow FT was visualized by confocal microscopy before and after cobalt quenching. After slow FT, calcein fluorescence was distributed both inside and outside GVs, regardless of lipid composition, indicating that calcein was encapsulated while equilibrating with the external solution during FT. Cobalt ions were then added to quench externally accessible calcein and assess membrane integrity of FT-generated GV. Although POPC produced only a small number of GVs, most POPC GVs retained intravesicular calcein fluorescence after cobalt addition, indicating that they formed relatively intact membrane compartments. In contrast, many PLPC GVs lost their encapsulated calcein signal after cobalt addition, suggesting that their membranes remained permeable to $\text{Co}^{2+}$ after FT. DOPC GVs frequently retained intravesicular calcein fluorescence after cobalt addition, although a subset of vesicles lost fluorescence, indicating that DOPC formed enlarged and relatively more intact compartments than PLPC. Scale bar: 10 $\mu\text{m}$. (b) To quantify the differences in calcein retention observed in (a), GV area and total intravesicular calcein fluorescence after $\text{Co}^{2+}$ quenching were analyzed for individual GVs. When membrane structures could not be separated because of apparent aggregation, the connected structure was analyzed as a single GV. POPC produced predominantly small GVs, whereas PLPC and DOPC generated substantially larger compartments. Among the latter two lipids, calcein fluorescence retained per unit GV area was higher for DOPC than for PLPC. Thus, whereas PLPC showed poor solute retention despite substantial GV growth, DOPC combined substantial vesicle growth with relatively high membrane integrity after slow FT.

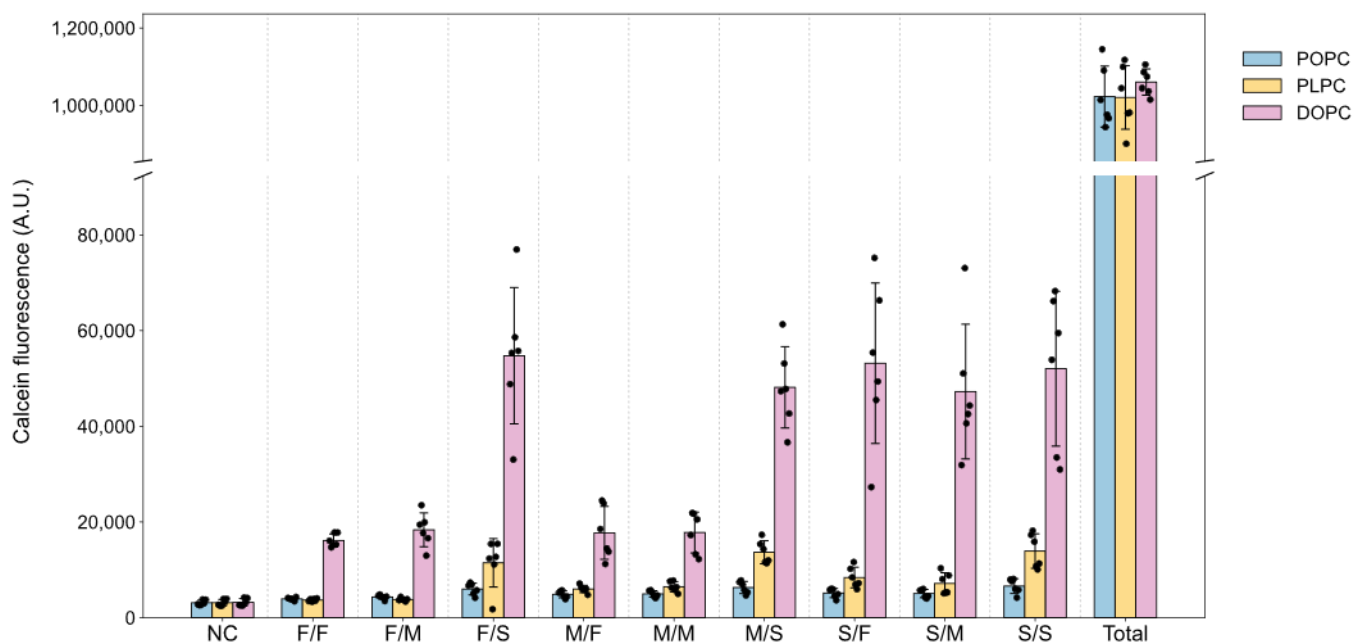

**Figure S6. Raw calcein fluorescence data used to calculate encapsulation efficiencies.** POPC, PLPC and DOPC empty LUVs were mixed with 10 $\mu\text{M}$ calcein and subjected to the indicated FT conditions. $\text{CoCl}_2$ was then added to a final concentration of 0.667 $\mu\text{M}$ to quench external calcein, and the remaining membrane-protected fluorescence was measured. NC denotes the calcein fluorescence intensity measured after $\text{CoCl}_2$ addition without FT treatment. Because calcein is impermeable to intact lipid membranes, it was not expected to enter the initial empty LUVs, and quenching of the external calcein reduced the fluorescence to near-background levels. Total denotes the fluorescence intensity of the initial mixture of 10 $\mu\text{M}$ calcein and LUVs measured without $\text{CoCl}_2$ quenching. Data are mean ± s.d. ($n = 6$).

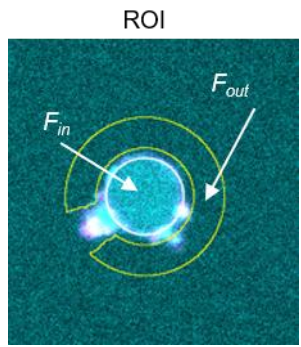

**Figure S7. Calculation of enrichment factors.** Enrichment factors ($EFs$) were calculated as $EF = F_{in} / F_{out}$, where $F_{in}$ is the mean fluorescence intensity inside each vesicle and $F_{out}$ is the mean fluorescence intensity in the surrounding external solution. Before quantification, a background image, obtained by averaging four images of FT samples without fluorescent solute, was subtracted pixel-wise from each fluorescence image. Vesicle interiors were manually defined as regions of interest (ROIs, outlined in white). For each vesicle, $F_{out}$ was measured from a 3-pixel-wide ring-shaped region positioned 1 pixel outside the vesicle boundary (outlined in yellow). When neighboring vesicles, aggregates or other fluorescent structures overlapped with the external ring-shaped region, the corresponding areas were excluded from the analysis.

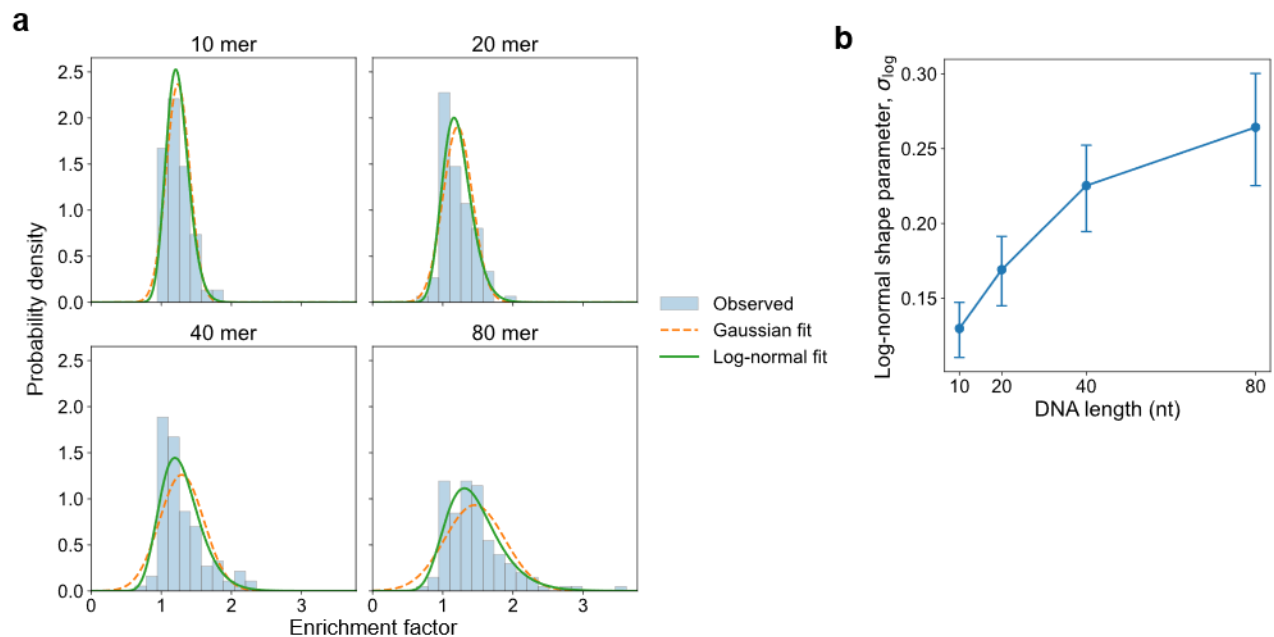

**Figure S8. Statistical analysis of enrichment factor distributions.** (a) Histograms of enrichment factor ($EF$) values for each DNA length, overlaid with Gaussian and log-normal fits. When molecular encapsulation is relatively uniform, $EF$ values may follow an approximately Gaussian distribution. By contrast, a log-normal distribution can arise when enrichment is influenced by multiple variable processes acting together, producing an asymmetric distribution with a long tail towards high $EF$ values. The right-skewed $EF$ distributions were better described by a log-normal than by a Gaussian model (See Table S2), suggesting that vesicle enrichment reflects the combined effects of processes such as freeze concentration, encapsulation and molecular retention, thereby generating greater heterogeneity than expected from simple encapsulation alone. (b) Dependence of the log-normal shape parameter, $\sigma$, on DNA length. In a log-normal distribution, $\sigma$ quantifies the spread of the distribution in log space; larger $\sigma$ values indicate broader heterogeneity and a heavier right tail. The increase in $\sigma$ with DNA length therefore indicates that enrichment became progressively more heterogeneous for longer DNA molecules, with a greater tendency for rare vesicles to exhibit exceptionally high $EF$ values. Error bars indicate 95% confidence intervals obtained by bootstrap resampling (10,000 iterations).

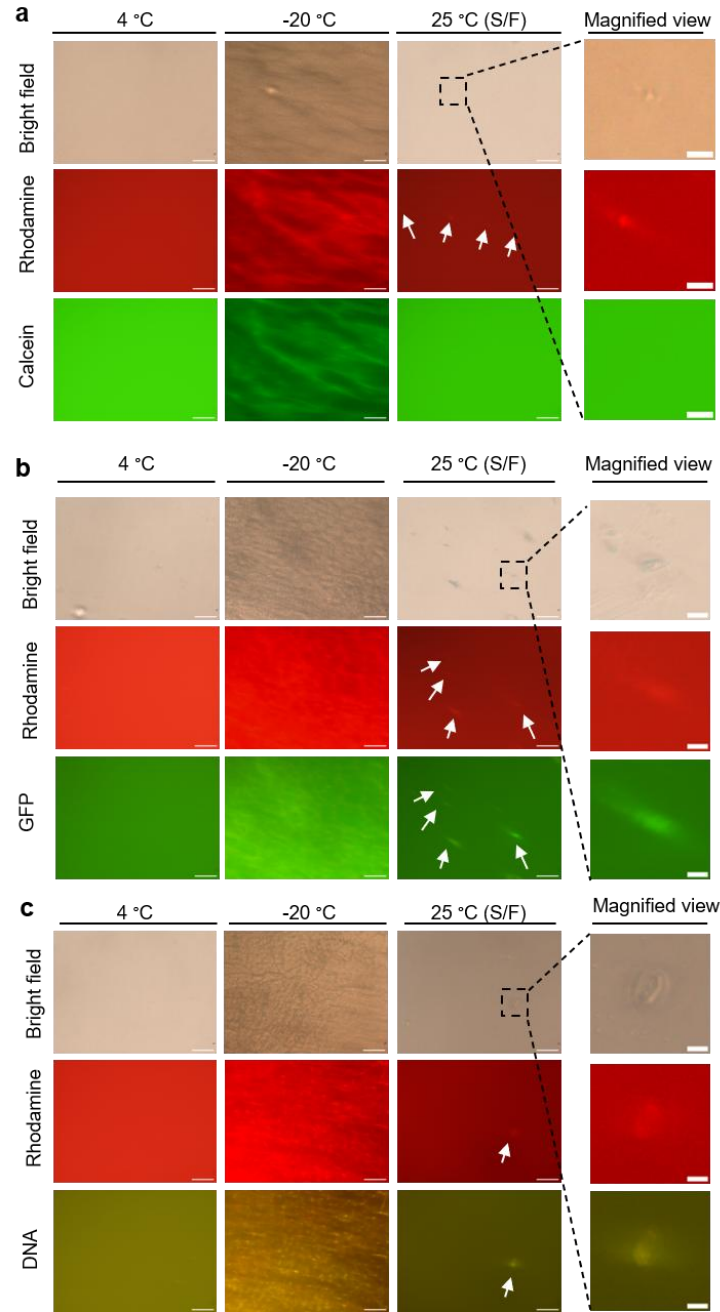

**Figure S9. *In situ* bright-field and fluorescence imaging of solute concentration during FT.** Rhodamine-labelled vesicles were mixed with (a) 10 μM calcein, (b) 1 μM GFP or (c) 10 nM DNA stained with 0.2× SYBR Gold, and imaged *in situ* before FT at 4 °C, in the frozen state after slow cooling to -20 °C (identical to the images shown in Fig. 3c) and after thawing (25 °C). For all three solutes, regions of high fluorescence intensity appeared upon ice formation and overlapped with dark interstitial regions between ice crystals in bright-field images. Rhodamine-labelled vesicles also accumulated in these regions, indicating that both solutes and lipid vesicles were freeze-concentrated within ice interstices (Fig. S10). After thawing, GFP- (b) and DNA-rich regions (c) remained partially colocalized with rhodamine-rich micrometer-sized structures as indicated by arrows (see magnified view), suggesting that concentrated biomolecules were retained within or associated with membrane-rich compartments. In contrast, calcein fluorescence became more uniformly distributed after thawing and showed little apparent colocalization with rhodamine-rich structures (a). Scale bars, 100 μm; magnified views, 20 μm.

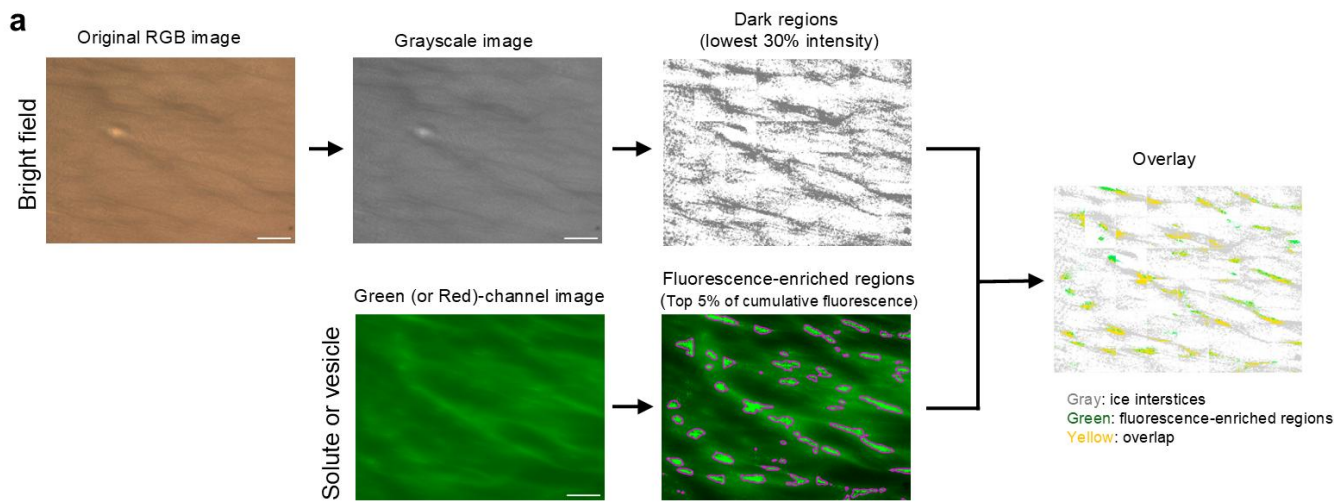

**b**

#### Definitions of dark regions

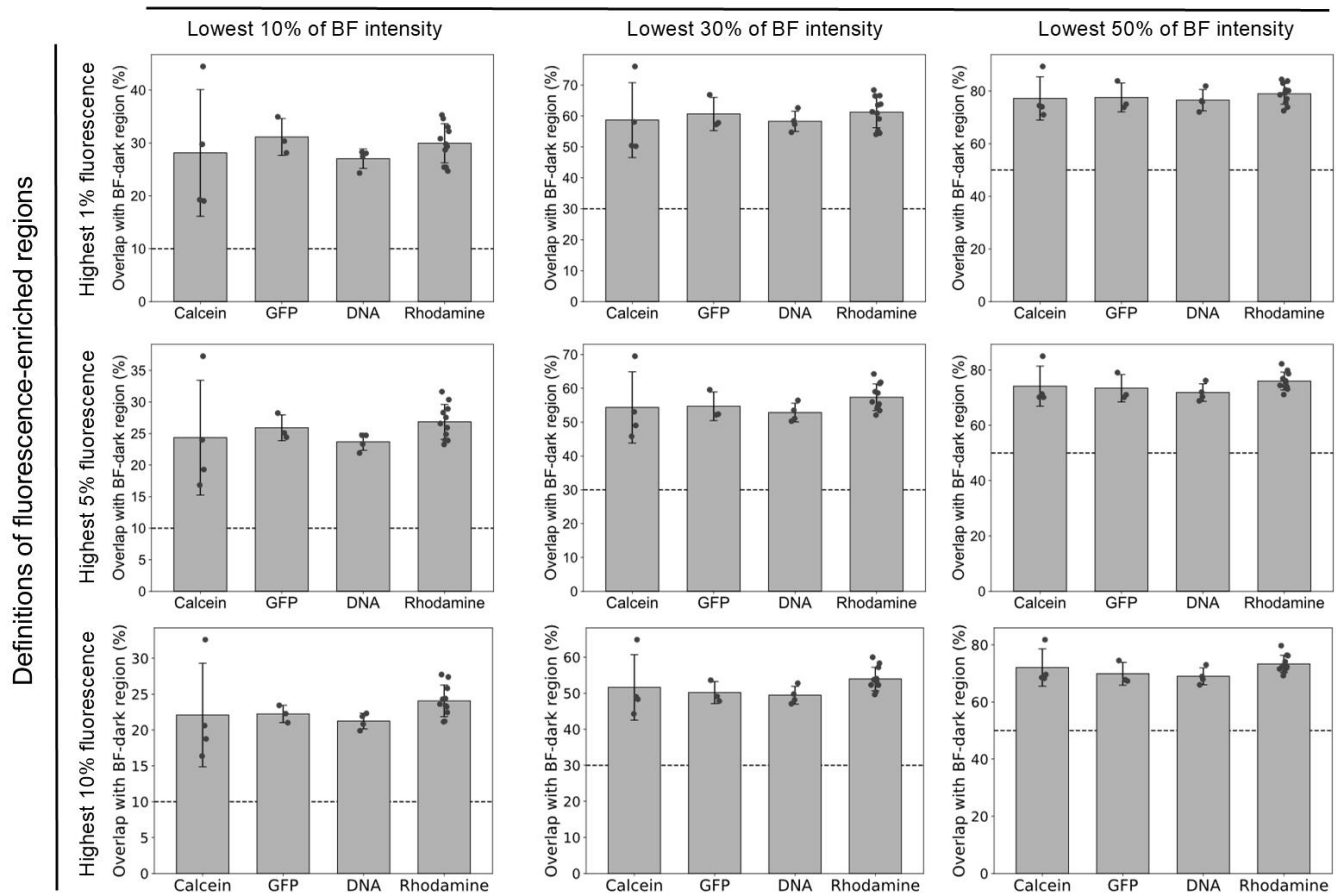

**Figure S10. Quantification of the spatial overlap between solute- or vesicle-enriched regions and ice interstices.**

(a) Image-analysis procedure used to quantify spatial overlap between fluorescence-enriched regions and ice interstices. To reduce the effect of uneven illumination across the field of view, each image was divided into $6 \times 8$ subregions and analyzed locally. Bright-field RGB images were converted to grayscale, and, as an example, pixels within the lowest 30% of bright-field intensity were defined as dark ice-interstitial regions. Fluorescence-rich regions were defined as the regions accounting for the upper cumulative 5% of the total fluorescence signal. The two regions were then overlaid to quantify their spatial overlap. In the example shown, gray indicates ice interstices, green indicates calcein-rich regions and yellow indicates their overlap. The overlap ratio was calculated as the area of the yellow region divided by the total area of the calcein-rich region (yellow + green). The same analysis was applied to GFP, DNA and rhodamine-labelled vesicles, with the resulting overlap ratios shown in Fig. 3d of the main text. If fluorescence-rich regions were randomly distributed, the expected overlap would be 30%, because 30% of the field was defined as interstitial. The measured overlap ratios were substantially higher, reaching $54.4 \pm 10.5\%$ for calcein ($n=4$), $54.7 \pm 4.22\%$ for GFP ($n=3$), $52.8 \pm 2.77\%$ for DNA ($n=4$), and $57.4 \pm 3.92\%$ for rhodamine-labeled vesicles ($n=11$) (mean ± s.d.). These results indicate preferential localization of solute-enriched regions within the interstitial regions between ice crystals. (b) Robustness of the spatial-overlap analysis to the percentile thresholds used to define fluorescence-enriched and dark bright-field regions. Fluorescence-enriched regions were defined using the highest 1%, 5% or 10% of fluorescence intensity, while dark bright-field regions were defined using the lowest 10%, 30% or 50% of bright-field intensity. Preferential localization of fluorescence-enriched regions within dark bright-field regions was consistently observed across all nine combinations of thresholds. Dashed lines indicate the overlap expected for random localization, corresponding to the fraction of the image defined as the dark bright-field region.

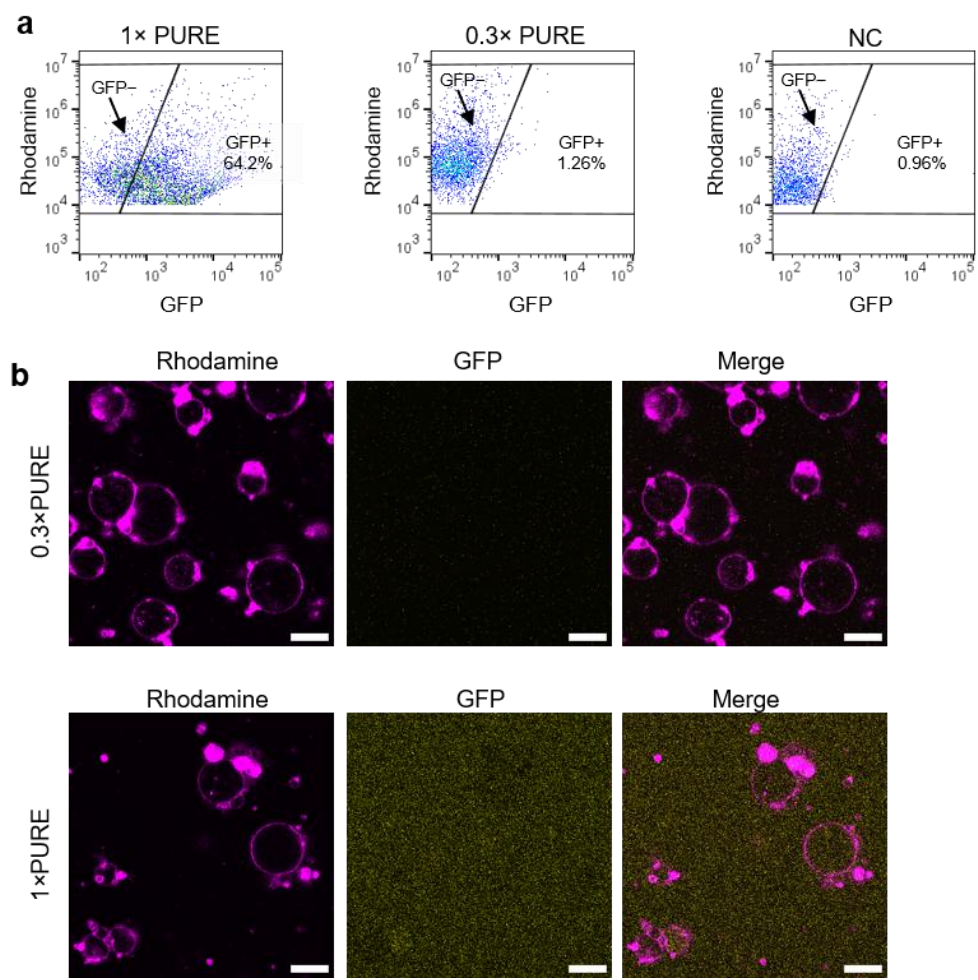

**Figure S11. PURE system activity in GUVs prepared by the emulsion-transfer method.** DOPC GUVs were prepared by the emulsion-transfer method<sup>3,4</sup> using PURE system solutions at either 0.3× or 1× the standard concentration as both the inner and outer aqueous phases. The inner and outer solutions were supplemented with 400 mM sucrose and 400 mM glucose, respectively, to provide a density contrast required for emulsion transfer. Because this method encapsulates the inner aqueous phase without an intrinsic molecular enrichment step, PURE components are expected to be encapsulated at approximately their supplied concentrations. (a) Flow-cytometric analysis of GUVs prepared with 1× or 0.3× PURE was performed using a CytoFLEX S flow cytometer (Beckman Coulter). During acquisition, thresholds of 500 were applied to both FSC and SSC. Detector gains were set to 4 for FSC, 12 for SSC, 1 for the GFP channel and 1 for the rhodamine channel. A total of 10,000 events were acquired and analyzed using FlowJo (v10.10.0; BD Biosciences). During analysis, events with rhodamine fluorescence intensities greater than $10^4$ were selected to exclude low-fluorescence small-particle and background events. Within this population, a negative control (NC) lacking GFP-encoding DNA was used to define the GFP-positive gate. GFP-positive GUVs accounted for 64.2% of the population at 1× PURE but only 1.26% at 0.3× PURE, indicating that 0.3× PURE supported little detectable GFP production after compartmentalization alone. Not all GUVs prepared with 1× PURE showed detectable GFP production, possibly owing to stochastic variation in the encapsulated concentrations of the ~70 PURE components, membrane-associated inhibition or effects of the sucrose on reaction efficiency or GFP fluorescence. (b) Representative confocal fluorescence images of GUVs prepared with 0.3× or 1× PURE. At 0.3× PURE, no detectable GFP production was observed either inside the GUVs or in the external solution, consistent with the low activity of 0.3× PURE in bulk, while positive control 1× PURE produced GFP both inside and outside the GUVs. These results indicate that membrane compartmentalization alone is insufficient to restore the activity of diluted 0.3× PURE.

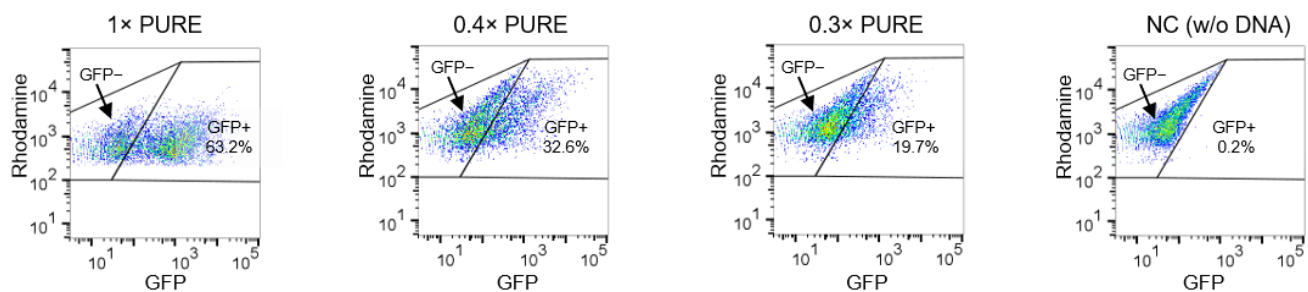

**Figure S12. Flow cytometric analysis of slow FT-driven activation of diluted PURE systems in GVs.** Empty DOPC vesicles ($1 \text{ mg mL}^{-1}$) were mixed with GFP-encoding DNA and PURE system components at matched dilutions of $1\times$, $0.4\times$ or $0.3\times$, corresponding to DNA concentrations of 10, 4 and 3 nM, respectively, and then subjected to slow FT. The resulting GVs were analyzed by flow cytometry using a FACSaria III cell sorter. During acquisition, thresholds of 500 were applied to both FSC and SSC. Detector voltages were set to 60 V for FSC, 340 V for SSC, 600 V for the GFP channel and 270 V for the rhodamine channel, and identical acquisition settings were used for all samples within the comparison. A total of 10,000 events were acquired for each sample. During subsequent analysis in FlowJo, events with rhodamine fluorescence intensities greater than 200 were selected to exclude low-fluorescence small-particle and background events. Within this population, a negative control (NC) containing $0.3\times$ PURE without GFP-encoding DNA was used to define the gates for GFP-negative and GFP-positive GVs. GFP-positive GVs were detected even from diluted PURE systems after slow FT. Within the GV population generated by slow FT, 63.2%, 32.6% and 19.7% of vesicles were GFP-positive for $1\times$, $0.4\times$ and $0.3\times$ PURE, respectively. After subtraction of the fluorescence measured in the NC, the mean GFP fluorescence intensities of GFP-positive GVs were 997, 469 and 353 a.u. for $1\times$, $0.4\times$ and $0.3\times$ PURE, respectively. To quantify GFP production at the population level, total GFP output was calculated by multiplying the number of GFP-positive GVs by their mean GFP fluorescence intensity. These results show that slow FT can generate membrane compartments supporting gene expression even when both the DNA template and PURE system are diluted to concentrations associated with reduced or undetectable activity in bulk solution.

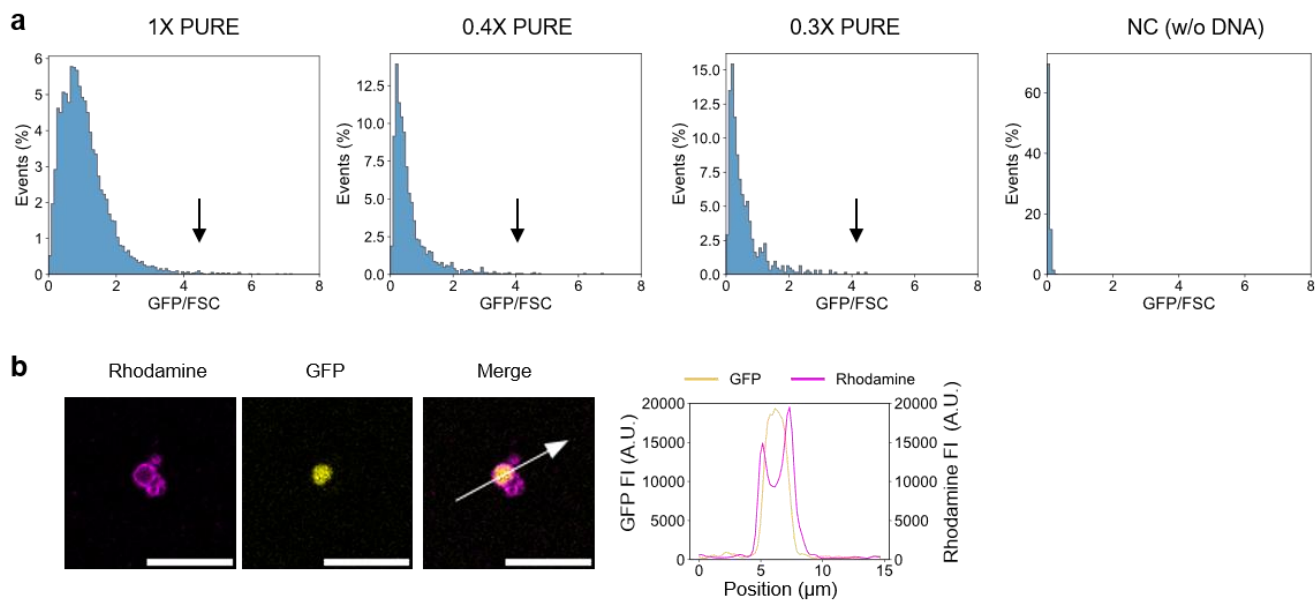

**Fig. S13. Highly GFP-expressing GV's generated by slow FT.** (a) Analysis of the flow cytometry data shown in Fig. S12. GFP/FSC was calculated for each individual GV as an approximate measure of GFP fluorescence relative to vesicle size. The distributions exhibited long tails extending toward high GFP/FSC values, indicating the presence of a small population of GV's with exceptionally high GFP fluorescence relative to their FSC signal. Arrows indicate the high-GFP/FSC region. (b) Representative confocal fluorescence image of a highly GFP-enriched GV generated by slow FT with 0.3× PURE. Scale bar: 10 $\mu\text{m}$.

**Table S1. Summary statistics for enrichment factor analysis.** For each solute, the table lists the molecular weight, molecular size (hydrodynamic diameter), number of analyzed GVs, and the mean and median enrichment factor (*EF*) values.

| Solutes | Molecular wight | molecular size | Sample number | Mean | Median |
| --- | --- | --- | --- | --- | --- |
| Calcein | 0.623 kDa | 1.48 nm<sup>a</sup> | 98 | 0.993 | 0.994 |
| Calcein NC |  |  | 89 | 0.198 | 0.181 |
| GFP | 48 kDa<br>(SNAP-GFP) | ~7.1 nm<sup>b</sup> | 73 | 7.497 | 6.425 |
| GFP NC |  |  | 68 | 0.286 | 0.279 |
| OA647 | 45 kDa | ~6.0 nm<sup>c</sup> | 78 | 2.023 | 1.757 |
| OA647 NC |  |  | 72 | 0.256 | 0.246 |
| DNA | 591 kDa<br>(GFP-encoding DNA, 956 bp) | n.d. | 152 | 2.830 | 1.622 |
| DNA NC |  |  | 110 | 0.149 | 0.137 |
| 10 mer ssDNA | 3.03 kDa | ~1.8 nm<sup>d</sup> | 95 | 1.238 | 1.212 |
| 20 mer ssDNA | 6.04 kDa | ~2.6 nm<sup>d</sup> | 95 | 1.212 | 1.158 |
| 40 mer ssDNA | 12.06 kDa | ~4.0 nm<sup>d</sup> | 118 | 1.291 | 1.188 |
| 80 mer ssDNA | 24.11 kDa | ~5.9 nm<sup>d</sup> | 128 | 1.455 | 1.395 |

<sup>a</sup> Hydrodynamic diameter of calcein was calculated as twice the reported Stokes–Einstein radius (0.74 nm)<sup>5</sup>.

<sup>b</sup> Hydrodynamic diameter of SNAP–GFP was approximated from the reported hydrodynamic size of a comparably sized GFP fusion protein<sup>6</sup>.

<sup>c</sup> Hydrodynamic diameter of OA647 was approximated from the reported hydrodynamic radius of native ovalbumin (2.98 nm); the contribution of Alexa Fluor 647 conjugation was not included<sup>7</sup>.

<sup>d</sup> Hydrodynamic diameters of ssDNA were estimated from published hydrodynamic-radius measurements of fluorescently labelled poly(dT) of corresponding lengths measured by fluorescence correlation spectroscopy<sup>8</sup>.

**Table S2. Comparison of distribution fits for enrichment factor values using the Akaike information criterion.** Enrichment factor (*EF*) distributions for 10-, 20-, 40- and 80 mer ssDNA were fitted with Gaussian, gamma, Weibull and log-normal distributions. These distributions were selected to compare a symmetric Gaussian model with three commonly used positive continuous distributions that can accommodate different degrees of right skewness and tail behavior. Model performance was evaluated using the Akaike information criterion (AIC)<sup>9</sup>, which accounts for both goodness of fit and model complexity. A lower AIC indicates greater relative support among the models tested. The log-normal distribution gave the lowest AIC for all four DNA lengths, indicating that it provided the best-supported fit among the candidate distributions.

| **DNA length** | **Gaussian** | **Gamma** | **Weibull** | **Log-normal** |
| --- | --- | --- | --- | --- |
| **10** | -64.64 | -72.35 | -43.45 | -75.49 |
| **20** | -23.49 | -28.85 | -11.88 | -30.29 |
| **40** | 67.37 | 48.23 | 79.07 | 41.18 |
| **80** | 149.96 | 121.94 | 159.08 | 113.17 |

### References

1. Song, L. Structure staphylococcal alpha-hemolysin, heptamerictransmembrane pore. *Science* **274**, 1859–1866 (1996).
2. Chiba, M., Miyazaki, M. & Ishiwata, S. Quantitative analysis of the lamellarity of giant liposomes prepared by the inverted emulsion method. *Biophys. J.* **107**, 346–354 (2014).
3. Pautot, S., Frisken, B. J. & Weitz, D. A. Production of unilamellar vesicles using an inverted emulsion. *Langmuir* **19**, 2870–2879 (2003).
4. Nishimura, K., Tsuru, S., Suzuki, H. & Yomo, T. Stochasticity in gene expression in a cell-sized compartment. *ACS Synth. Biol.* **4**, 566–576 (2015).
5. Tamba, Y., Ariyama, H., Levadny, V. & Yamazaki, M. Kinetic pathway of antimicrobial peptide magainin 2-induced pore formation in lipid membranes. *J. Phys. Chem. B* **114**, 12018–12026 (2010).
6. Hink, M. A. *et al.* Structural dynamics of green fluorescent protein alone and fused with a single chain Fv protein. *J. Biol. Chem.* **275**, 17556–17560 (2000).
7. Smilgies, D.-M. & Folta-Stogniew, E. Molecular weight-gyration radius relation of globular proteins: a comparison of light scattering, small-angle X-ray scattering and structure-based data. *J. Appl. Crystallogr.* **48**, 1604–1606 (2015).
8. Doose, S., Barsch, H. & Sauer, M. Polymer properties of polythymine as revealed by translational diffusion. *Biophys. J.* **93**, 1224–1234 (2007).
9. Akaike, H. A new look at the statistical model identification. *IEEE Trans. Automat. Contr.* **19**, 716–723 (1974).
